# Sex-lethal is recruited to chromatin to promote neuronal tRNA synthesis in males through RNA Polymerase III regulation

**DOI:** 10.1101/2025.04.25.650657

**Authors:** Freya Storer, Colin D. McClure, Alicia Estacio Gomez, Tsz Lam Wong, Lucy J. Minkley, Tony D. Southall

**Affiliations:** Department of Life Sciences, Imperial College London, Sir Ernst Chain Building, London, SW7 2AZ, UK

## Abstract

The RNA-binding protein Sex-lethal (Sxl) is classically known as a master regulator of sex determination and mRNA splicing in *Drosophila melanogaster*. However, this role is not conserved across species, and functions beyond this canonical pathway remain poorly understood. In this study, we uncover a splicing-independent role for Sxl at the chromatin level in the *Drosophila* brain. Using Targeted DamID (TaDa) profiling in neurons, we identify widespread recruitment of Sxl to promoter regions, independent of sex or RNA binding activity. Notably, Sxl chromatin occupancy exhibits near-complete overlap with Polr3E (RPC37), an RNA Polymerase III subunit, with Sxl binding abolished upon *Polr3E* knockdown. Depletion of Sxl in mature male neurons induces widespread transcriptional changes, particularly in metabolic genes, and improves negative geotaxis during ageing, phenotypes that closely mirror *Polr3E* knockdown. Conversely, overexpression of the brain-specific *Sxl^RAC^* transcript leads to enhanced tRNA synthesis and upregulated metabolic gene expression. Together, these findings reveal a previously unrecognised role for Sxl in regulating Pol III activity via Polr3E, regulating tRNA synthesis and supporting neuronal metabolism. Given the emerging tie between Pol III regulation and neuronal ageing, our study highlights Sxl as a novel modulator of neuronal homeostasis.

## Introduction

Sexual dimorphism in *Drosophila melanogaster* is tightly controlled by one of the most established pathways in biology. At its core lies Sex lethal (Sxl), a splicing factor and master regulator that acts as a binary switch, controlling the sexual identity of the organism. In early female embryos, the ratio of X-linked signalling elements activates Sxl’s ‘early’ promoter, initiating expression of the *Sxl^early^* transcript (Sanchez *et al*., 1994). Sxl^early^ protein, once translated, directs the female-specific splicing of the *Sxl^late^* transcripts, which is constitutively transcribed (Keyes *et al*., 1992; Salz and Erickson, 2010). This autoregulatory splicing ensures the continued production of full-length, functional Sxl protein in females, maintaining its own expression and initiating the downstream female-specific sex determination cascade (Bell *et al*., 1991). In males, where *Sxl^early^* is not produced, *Sxl^late^* undergoes default splicing, generating a truncated protein with no known function. This binary mechanism ensures a stable and heritable cellular memory of sexual identity throughout development (Fig. 1a).

**Figure 1.**
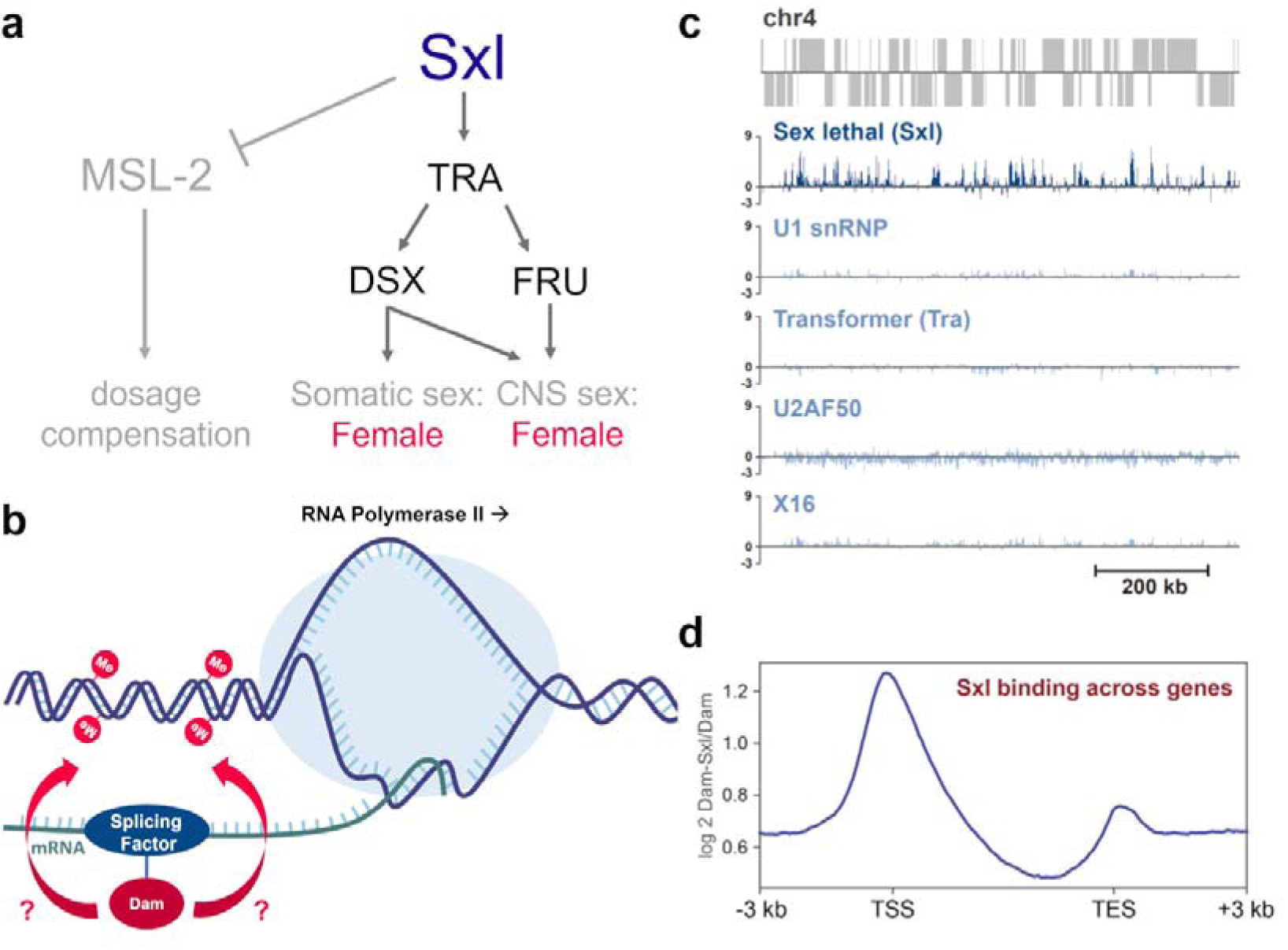
Sxl robustly associates with chromatin while other splicing factors do not. **a**, Overview of Sxl’s role in sex determination in *Drosophila melanogaster* (adapted from Robinett *et al*., 2010). **b**, Schematic outlining the use of Targeted DamID (TaDa) to assess whether splicing factors make transient contacts with chromatin. **c**, TaDa profiles for Sxl and four additional splicing factors across chromosome 4 in neurons of *Drosophila* larval brains. The *y*-axis represents the log_ ratio of Dam-factor over Dam-alone. **d**, Metagene plot showing average Sxl binding across gene bodies, aligned to transcription start sites (TSS) and transcription end sites (TES). Data underlying this figure are available in Table S1.

Although essential for sexual differentiation, the function of Sxl has long been considered limited to a number of well-defined roles. Its primary function is to regulate the female-specific splicing of *transformer (tra)* pre-mRNA, a key step in the sex determination hierarchy (Boggs *et al*., 1987). Sxl promotes exon skipping in *tra*, preventing premature translation termination and enabling the production of functional Tra protein. In turn, Tra drives feminisation of tissues through the alternative splicing of transcription factor genes *Doublesex* (*Dsx*) and *Fruitless* (*Fru*) by producing female-specific isoforms of each protein (Robinett *et al*., 2010). These factors drive transcriptional changes that lead to the majority of sexual dimorphic physiology and behaviour in *Drosophila*. In addition to its role in splicing regulation, Sxl inhibits translation of *Male-Sex-lethal 2* (*msl-2*), thereby preventing assembly of the dosage compensation complex (DCC), which otherwise upregulates X-linked gene expression in males (Bashaw and Baker, 1997; Conrad and Akhtar, 2012).

The full-length Sxl protein contains two RNA-Binding Domains (RBDs), each accommodating two highly conserved ribonucleoprotein (RNP) motifs that recognise poly(U) sequences to regulate splicing (Handa *et al*., 1999). In addition to RNA recognition, these domains mediate interactions with other proteins, including splicing factor Snf16 and the RNA polymerase III subunit E (Polr3E) (Dong and Bell, 1999). Beyond the RBDs, Sxl possesses a 120-residue N-terminal domain and an 80-residue C-terminal domain. The first 40 amino acids of the N-terminus are critical for cooperative RNA binding (Wang and Bell, 1996). Loss of this region uncouples Sxl’s sex determination and dosage compensation functions, whereby the protein retains its ability to repress *msl-2* translation but fails to promote female-specific splicing of *tra* (Yanowitz *et al*., 1999).

An intriguing aspect of Sxl biology is the expression of a near full-length isoform, Sxl^RAC^, in the male nervous system (Bopp *et al*., 1991; Cline *et al*., 2010; Moschall *et al*., 2018). Unlike the canonical female-specific isoforms, Sxl^RAC^ is expressed in both sexes but appears restricted to the nervous system male pupae and adults (Cline *et al*., 2010; Moschall *et al*., 2018). Notably, Sxl^RAC^ lacks the first 23 amino acids of the N-terminal region, an area critical for promoting female-specific splicing of *tra* (Wang and Bell, 1994; Yanowitz *et al*., 1999). Consistent with this truncation, full-length Tra protein is absent in males (Boggs *et al*., 1987), suggesting that Sxl^RAC^ functions independently of its canonical role in alternative splicing.

Despite extensive characterisation of Sxl as the master regulator of sex determination, emerging evidence points to additional, Tra-independent functions. A phenomenon termed Tra-insufficient Feminisation (TIF) highlights physiological and transcriptional effects that cannot be explained by the canonical Sxl-Tra-Dsx/Fru axis (Evans and Cline, 2013). TIF includes i) partial feminisation of *Sxl* mutant lines rescued by Tra2, ii) transcriptional changes independent of Dsx and Fru (Goldman and Arbeitman, 2007), and iii) phenotypes following *Sxl* knockdown that are not replicated by *tra* depletion (Sawala and Gould, 2017). Collectively, these findings support the existence of an alternative, yet unidentified, mode of Sxl action beyond its canonical role in sex-specific splicing.

## Results

### Sxl robustly associates with chromatin while other splicing factors do not

The majority of mRNA splicing occurs co-transcriptionally, bringing factors into close proximity with DNA (Herzel *et al*., 2017). To investigate whether splicing factors transiently associate with chromatin *in vivo*, we employed Targeted DamID (TaDa), a cell-type specific method for profiling protein-DNA interactions without the need for antibodies or cell isolation (Southall *et al*., 2013). In this system, the protein of interest is fused to the *E.coli* DNA adenine methyltransferase (Dam), which deposits methylation marks at nearby GATC motifs, enabling genome-wide mapping of protein occupancy. Towards this, splicing factors were tagged with Dam to assess whether this enabled profiling of their transient interactions with chromatin in neurons of the developing larval brain (Fig. 1b). Dam fusions to U1, Tra, U2AF50 and x16 yielded no significant chromatin-associated signal (Fig. 1c), suggesting limited or highly transient interactions. In contrast, Sxl exhibited robust chromatin binding, with marked enrichment at transcriptional start sites (TSS) (Fig. 1d). Notably, Sxl binding encompassed a substantial proportion of the genome, with occupancy detected at 53% of expressed genes (Fig. S1, Table S1). This widespread chromatin occupancy suggests that Sxl may have a broader role in transcriptional regulation beyond its established function in alternative splicing.

### Sxl associates with chromatin independently of sex and RNA binding ability

Full-length Sxl, which mediates alternative splicing of *tra* pre-mRNA, is expressed in females but absent in males. To determine whether sex-specific factors influence Sxl’s chromatin binding, we performed TaDa profiling of Sxl in both males and females. Interestingly, no significant differences in binding were observed between the sexes, suggesting a role besides sex determination (Fig. 2). To explore whether Sxl’s chromatin association was due to its interaction with RNA substrates, two mutant versions of Sxl were generated, incapable of binding RNA. The first mutant (*Sxl^GS^*) replaces the RNA recognition motifs (RNPs) with glycine/serine linkers (Handa *et al*., 1999), while the second mutant (*Sxl^RNA^*) carries mutations in key amino acids within the RNPs (Y168A, F170D, V254A, F256D) (Lisbin *et al*., 2000). Previous studies have shown that similar mutations in other RNA-binding proteins, such as *Drosophila* ELAV, abolish RNA binding (Lisbin *et al*., 2000). Next, the chromatin binding function of these mutants were assessed using TaDa. Strikingly, both the Sxl^GS^ and Sxl^RNA^ lines were shown to bind chromatin (Fig. 2), as did mutants lacking either the N- or C- terminal regions. These results suggest that a cofactor, likely interacting with Sxl’s central domain (containing RNA-binding domains), may mediate Sxl’s recruitment to chromatin and is unlikely linked to RNA binding function. A strong candidate for this interaction is the Pol III subunit Polr3E, previously shown to bind Sxl in a yeast two-hybrid screen (Dong and Bell, 1999).

**Figure 2.**
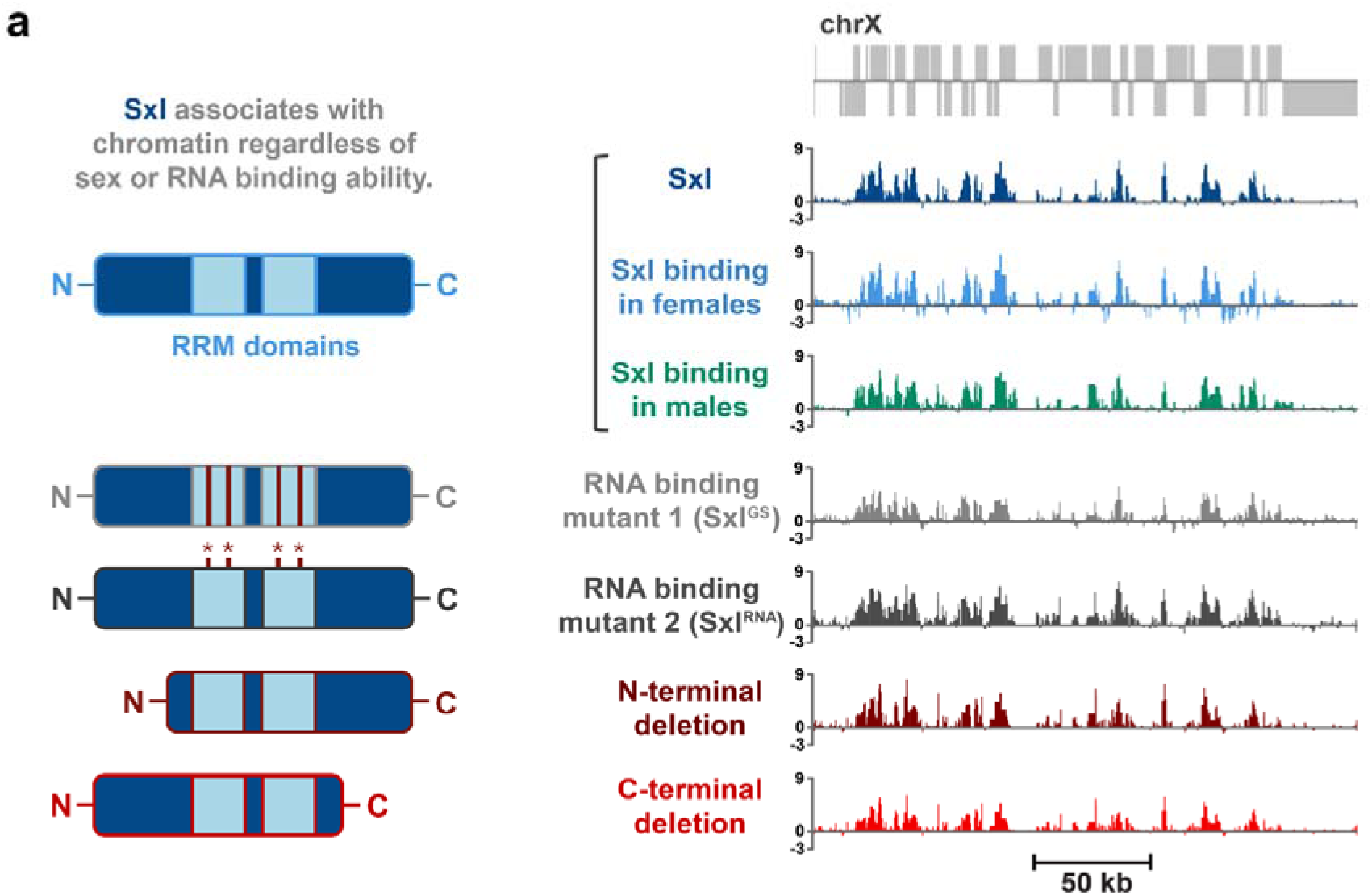
Sxl associates with chromatin independently of sex and its ability to bind RNA. **a**, Targeted DamID data showing chromatin occupancy for *wild type* Sxl in males and females, alongside Sxl variants with disrupted RNA-binding capacity. Pale blue regions denote RNA recognition motifs (RRMs). Red lines indicate positions where RNP sites were replaced with glycine-serine linkers. Asterisks mark individual point mutations (Y168A, F170D, V254A, F256D). Data underlying this figure are available in Tables S2, S3, S4 and S5.

### Polr3E is required for Sxl’s association with chromatin

To assess whether Polr3E mediates Sxl recruitment to chromatin, profiling of Polr3E binding across the genome in larval neurons was assessed using TaDa. Remarkably, it was found that Polr3E shared a highly similar binding profile with Sxl (Fig. 3a).

**Figure 3.**
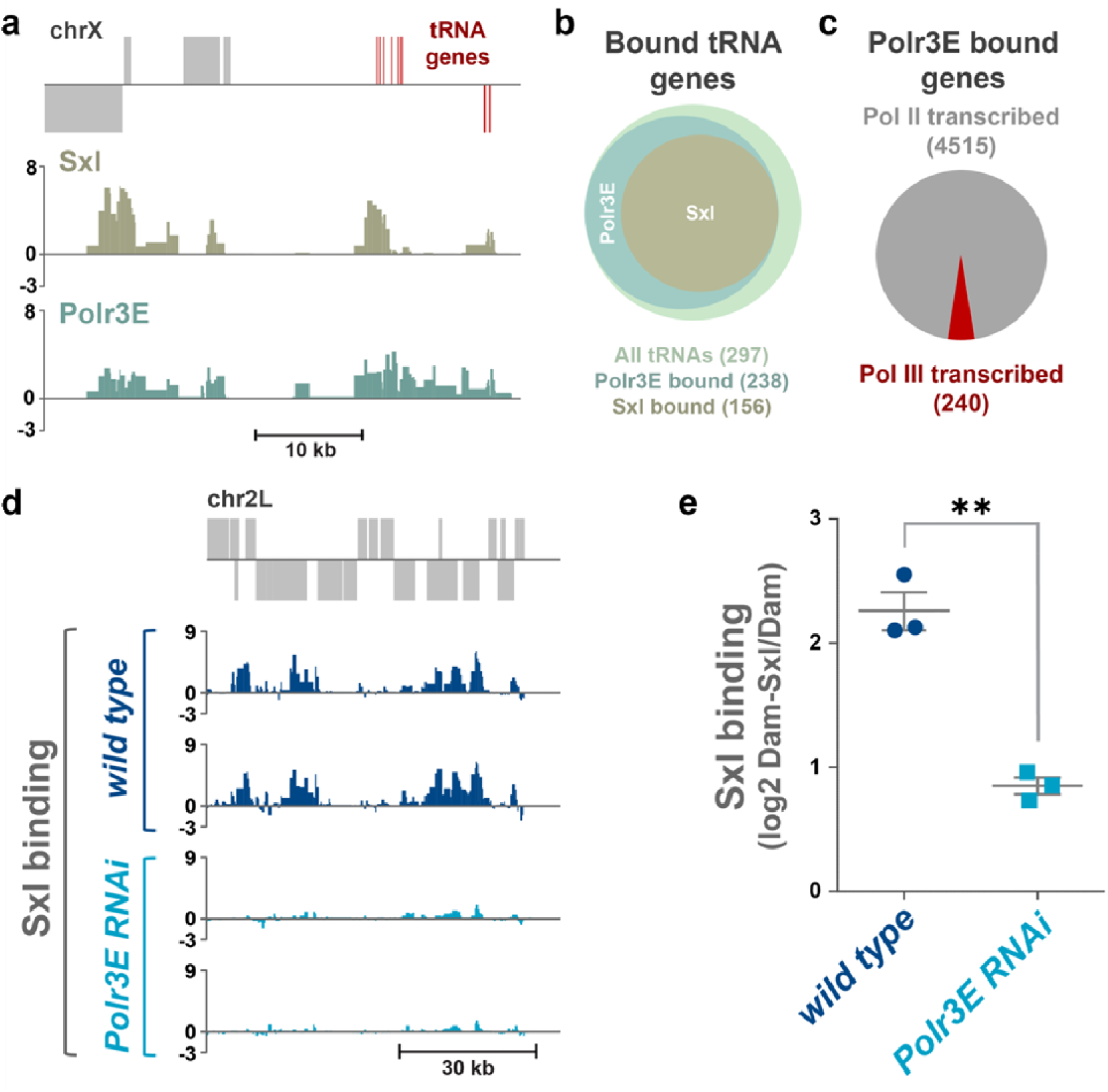
Polr3E is required for Sxl’s association with chromatin. **a**, Genome- wide binding of Polr3E shows strong overlap with Sxl chromatin occupancy using TaDa. **b,** Polr3E binds the majority of annotated tRNA genes, with Sxl binding a defined subset. **c,** Polr3E is recruited to both RNA polymerase II and RNA polymerase III-transcribed loci. **d,** Chromatin binding of Sxl is markedly reduced in neurons depleted of Polr3E by RNAi. **e,** Quantification reveals a significant reduction in Sxl binding upon *Polr3E* knockdown (*p*=0.006, unpaired *t*-test). Data underlying this figure are available in Tables S6. S7 and S8.

Both proteins bound to a significant number of Pol III-transcribed genes, including tRNA genes (Fig. 3b), but also a large subset of Pol II-transcribed genes (Fig. 3c). This observation is not unprecedented, as *Drosophila* TFIIC, a component of the Pol III transcriptional machinery, also associates with many Pol II-transcribed genes (∼2,400) (Van Bortle *et al*., 2014). To directly test whether Polr3E is required for Sxl binding, *Polr3E* was knocked down in larval neurons while simultaneously profiling Sxl Binding via TaDa. This resulted in a significant reduction of Sxl binding (*p=*0.0064; unpaired *t-*test; Fig. 3d-e, Table S8), confirming that Polr3E is essential for Sxl recruitment to chromatin.

### *Sxl* knockdown phenocopies *Polr3E* Loss in adult males

In order to further investigate Sxl’s role in the *Drosophila* nervous system, its expression and functional impact was examined in adult male neurons. This approach enabled the isolation of sex-independent functions, avoiding confounding effects related to its canonical role in female sex determination. Using a CRISPR knock-in line (*Sxl-T2A-GAL4*) (Moschal *et al*., 2018), we observed that Sxl expression is broadly distributed throughout the adult male brain, in contrast to its more limited pattern during larval stages, and is restricted to neurons (Fig. S2). To assess functional consequences, *Sxl* was knocked down in adult neurons using *nSyb-*GAL4, bypassing developmental effects using *tub-*GAL80^ts^. Depletion resulted in a mild but consistent reduction in survival (*p*<0.05, Gehan-Breslow-Wilcoxon test; Fig. 4a, Fig. S3) and age-related improvement of climbing ability after 21 days (*p*<0.05, two-way ANOVA; Fig. 4b), suggesting a role in neuronal maintenance.

**Figure 4.**
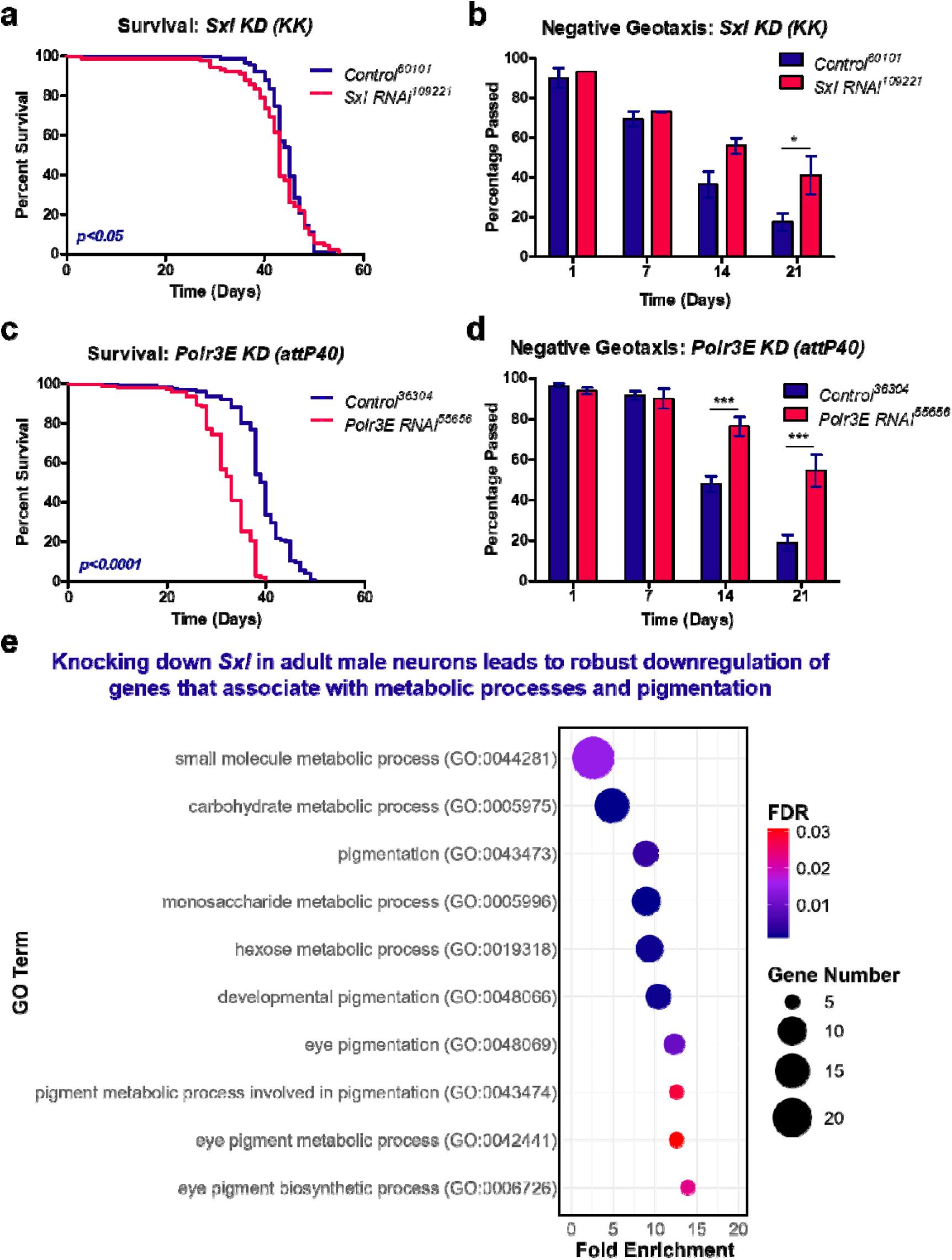
Sxl knockdown phenocopies Polr3E Loss in adult males. **a**, Survival curves of adult males following pan-neuronal knockdown of *Sxl* (VDRC #109221), compared to RNAi controls (VDRC #60101) (n>90 per group). Statistical significance assessed using Gehan–Breslow–Wilcoxon test. **b,** Negative geotaxis assay showing improved climbing ability following *Sxl* knockdown in male neurons. *y*-axis indicates the percentage of flies surpassing the 4cm midpoint. Data analysed by two-way ANOVA with Bonferroni post hoc test; \**p*<0.05. **c,** Survival curves following neuronal knockdown of *Polr3E* (BDSC # 55656) compared to RNAi controls (BDSC #36304) (n > 120 per group), mirroring the phenotype observed with *Sxl* depletion. Statistical significance assessed using Gehan–Breslow–Wilcoxon test. **d,** Improved climbing performance in *Polr3E* knockdown flies after day 14 post-eclosion; \*\*\**p* < 0.001. **e,** Gene ontology enrichment analysis of transcripts downregulated after *Sxl* knockdown in adult male neurons. Dot plot shows the top 10 GO terms ranked by fold enrichment (*x*-axis); dot size indicates the number of gene hits, and color reflects FDR. Data underlying this figure are available in Tables S9, S10, S11, S12 and S13.

Strikingly, knockdown of *Polr3E* produced comparable phenotypes, including reduced survival (*p*<0.0001, Gehan-Breslow-Wilcoxon test; Fig. 4c) and climbing improvement after day 14 (*p*<0.001, two-way ANOVA; Fig. 4d). Given previous evidence linking Pol III modulation to age-related phenotypes (Ureña *et al*., 2024; Malik *et al*., 2024), these findings point to a potential role for Sxl in similar pathways. To explore this further, RNA-seq was performed in young males (4 DPE) following *Sxl* knockdown in neurons and observed significant downregulation of genes involved in metabolic processes (Fig. 4e). *Polr3E* knockdown mirrored transcriptional changes, with a strong positive correlation (Spearman *r*=0.396, *p*<0.0001) between the two datasets (Fig. S3). Crucially, when exploring transcriptional changes of Dam- Sxl targets, it was noted that over 69% of targets were significantly downregulated in *Polr3E* knockdowns, reflecting over 65% targets downregulated in *Sxl* knockdowns (Fig. S3). Together, these results reveal a novel role for Sxl in regulating neuronal metabolism and a likely cooperation with Polr3E.

### Elevated Sxl^RAC^ leads to changes in tRNA synthesis

To investigate the function of the head-specific isoform Sxl^RAC^ (Cline *et al*., 2010), a *UAS-Sxl^RAC^* transgenic line was generated to assess its effects in mature male neurons. Overexpression of *Sxl^RAC^* significantly reduced survival relative to controls (*p*<0.0001, Gehan-Breslow-Wilcoxon test; Fig. 5a) and impaired climbing ability with age (*p*<0.01, two-way ANOVA; Fig. 5b), in contrast to the phenotypes observed upon gene knockdown. RNAseq analysis revealed widespread transcriptional changes in *Sxl^RAC^*-expressing neurons, including significant upregulation of genes associated with metabolism and translation (Fig. S4). Expression of *UAS-Sxl^RNA^*, with no RNA binding capacity recapitulated these phenotypes (Fig. S5), suggesting this function is not required in this context. Equally, ectopic expression of both *Sxl^RNA^* in *elav*-GAL4 (neuronal) and *tub*-GAL4 (ubiquitous) backgrounds did not induce lethality in males, confirming loss of RNA-binding function (Table S34). Given the phenotypic convergence between *Sxl* and *Polr3E* perturbation (Fig. 4) and their shared chromatin localisation at tRNA loci (Fig. 3b), we examined the impact of elevated Sxl^RAC^ on tRNA abundance. Small RNA sequencing revealed altered tRNA (Fig. 5c) and other small RNA profiles (Fig. 5d) in *Sxl^RAC^*-expressing males, with 72% of significantly affected tRNAs upregulated (Fig. 5e), including a striking enrichment for Lys-tRNAs (Table S16). Precursor tRNA levels, including *pre-tRNA^His^*, *pre-tRNA^TAT^*and *pre-tRNA^CAC^*, were elevated relative to the Pol II–specific U3 transcript, as measured by qPCR (Fig. S5), with reversed trends in *Sxl* knockdowns (Tables S40 and S41). To assess any effect translational output, O-propargyl-puromycin (OPP) labeling was employed to track nascent polypeptides. *mCherry* expression under *Sxl-T2A-GAL4* control revealed high Sxl levels in the mushroom bodies and medulla (Fig. S6), with Sxl intensity strongly correlating with OPP signal across brain regions (*R*coloc=0.81; Fig. 5f). Together, these results suggest a functional link between Sxl expression, tRNA synthesis, and translational activity in neurons.

**Figure 5.**
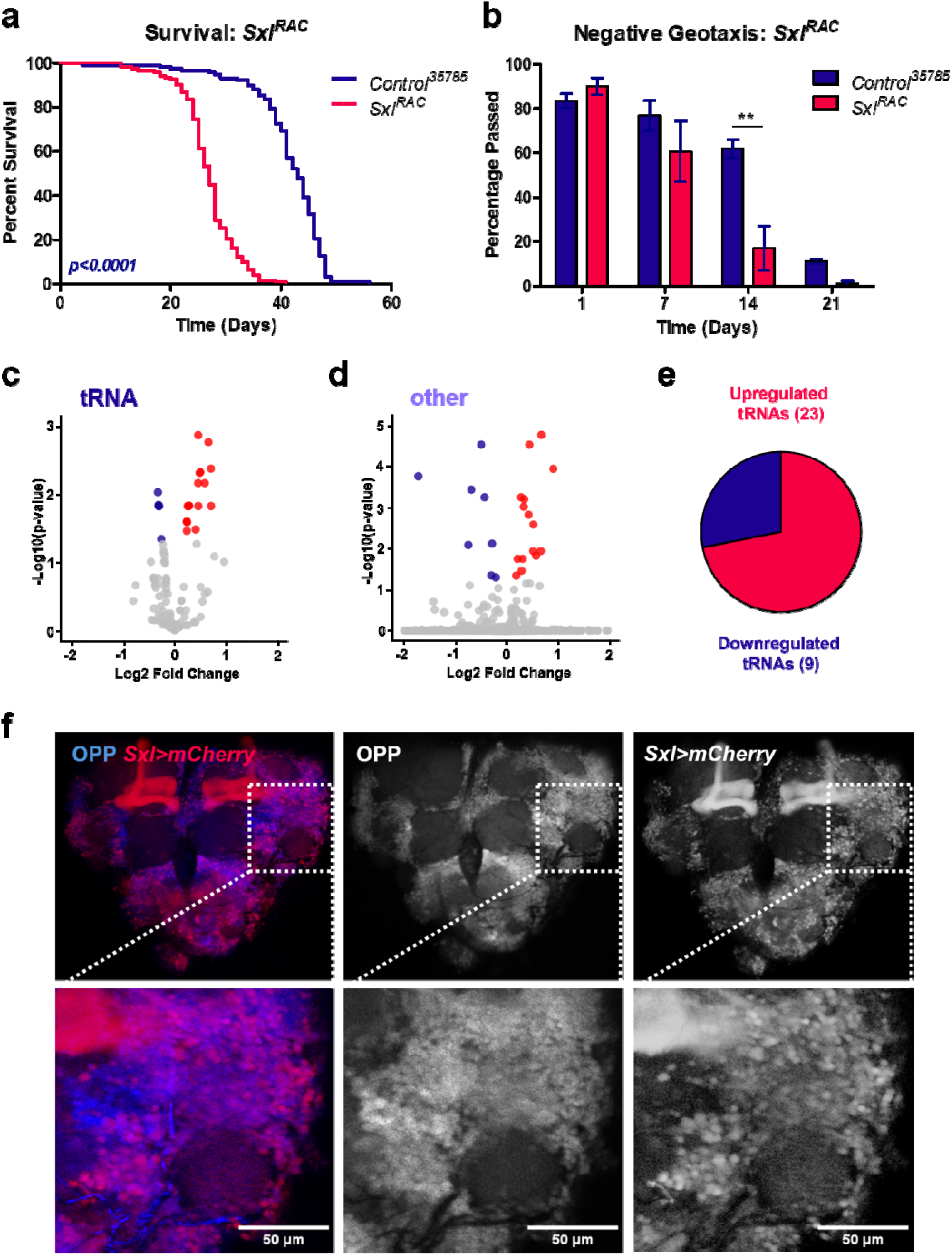
Elevated Sxl^RAC^ leads to changes in tRNA synthesis. **a**, Survival curves of adult males following pan-neuronal expression of *Sxl^RAC^*, compared to *mCherry* controls (n>119 per group). Statistical significance assessed using Gehan–Breslow–Wilcoxon test. **b,** Negative geotaxis assay showing reduced climbing ability following *Sxl^RAC^* overexpression in male neurons. *y*-axis represents the percentage of flies surpassing the 4cm midpoint. Data were analysed by two-way ANOVA with Bonferroni post hoc correction; \*\**p*<0.01. **c,** Volcano plot showing differential expression of tRNAs in adult male heads comparing *Sxl^RAC^*-expressing samples compared to controls. Significantly upregulated tRNAs (*p<*0.05) are highlighted in red; downregulated tRNAs (*p<*0.05) in blue. **d,** “Other” indicates additional small RNA species, including rRNA, miRNA, ncRNA, snoRNA and snRNA. **e,** Pie chart illustrating that the majority (72%) changes in tRNA levels due to upregulation. **f,** Immunolabelled sections of the adult male brain showing protein synthesis (OPP, blue) and *Sxl* expression (mCherry, red). Images captured at 20x magnification; scale bar, 50μm. Colocalisation analysis yielded *R*coloc=0.81. Data underlying this figure are available in Tables S14, S15, S16, S17 and S18.

## Discussion

Sex-lethal (Sxl), a master regulator of sex determination in the fruit fly, is known to complete this function through RNA splicing. Our study uncovers a previously unrecognised role for Sxl beyond this well-studied mechanism. Using Targeted DamID, we demonstrate that Sxl is robustly recruited to chromatin in the *Drosophila* brain, independent of RNA-binding and sex, and this recruitment is mediated through its interaction with Polr3E. Functional analyses reveal that Sxl influences Pol III-driven transcription, particularly of tRNAs, and plays an important role in maintaining neuronal health. Knockdown and overexpression studies in adult male neurons expose novel, sex-independent functions for Sxl, including regulation of metabolic gene expression, lifespan, and age-related motor performance. Together, these findings establish Sxl as a chromatin-associated modulator of Pol III activity and neuronal homeostasis.

### Sxl as a chromatin interactor

A key discovery from our research is the identification of Sxl’s chromatin association, with enriched binding at promoters and transcriptional start sites (Fig. 1). Notably, this chromatin recruitment occurs independently of Sxl’s splicing activity and RNA-binding ability and is evident in both male and female animals (Fig. 2).

Our studies further suggest that Sxl is widely expressed during late-stage brain development, with distinct and context-dependent expression patterns in adult male tissue, implying a requirement for functionally specialised roles. Mutant analysis confirms that chromatin recruitment is independent of both the N- and C-terminus of Sxl (Fig. 2), reinforcing its splicing-independent role and highlighting the importance of its RNA-binding domain, which likely mediates its interactions but not through RNA binding *per se*. These findings open new avenues for investigating the role of Sxl and other RNA-binding proteins moonlighting as chromatin interactors (Rafiee *et al*., 2021; Ray *et al*., 2023).

### RNA Polymerase III is modulated by Sxl

In the case of Sxl, RBD-1 is responsible for its interaction with binding partners like Polr3E (Dong and Bell, 1999). The Targeted DamID data presented here reveals a striking co-occupancy of Sxl and Polr3E, with Sxl binding enriched at tRNA loci as well as Pol II-specific genes (Fig. 3). Indeed, knockdown of *Polr3E* leads to a loss of Sxl binding to chromatin, suggesting a cooperative mechanism. We speculate that this interaction not only facilitates Pol III transcription but may also influence chromatin architecture and RNA Pol II-driven transcription as observed with Pol III regulation in other organisms (Carriere *et al*., 2011; Wang *et al*., 2014; Jiang *et al*., 2022). Although direct evidence of *Polr3E* function in *Drosophila* neurons is limited, recent studies indicate that inhibition of Pol III activity yields phenotypes analogous to those observed in our functional knockdown experiments (Malik *et al*., 2024; Ureña *et al*., 2024). In particular, knockdown of *Polr3E* in adult male neurons results in a modest reduction in survival, but significantly enhances performance in the negative geotaxis assay as the flies age (Fig. 4). This phenotype mirrors the effects seen in Sxl knockdown experiments, supporting the hypothesis that Sxl and *Polr3E* work together to maintain neuronal function. Transcriptional analyses further reinforce this, with correlation tests revealing a significant relationship between the two datasets (Fig. S3) and highlighting metabolism as a key downstream process.

RNA Pol III is expressed ubiquitously during development and in a cell-selective manner in adulthood, where its activity must be tightly regulated (Takada *et al*., 2000; Rideout *et al*., 2011). Traditionally, RNA Pol III has been understood as a core enzyme responsible for transcribing small, essential non-coding RNAs like tRNAs, 5S rRNA, and U6 snRNA, primarily to support basal cellular function. However, recent evidence shows that Pol III activity is often elevated or specialized, reflecting the unique metabolic and transcriptional needs of highly active cells, particularly neurons (Marshall *et al*., 2012; Ishimura *et al*., 2014). This likely explains Sxl’s role in the *Drosophila* nervous system; to regulate Pol III activity in late brain development through its interaction with Polr3E. Based on its specific expression patterns, we hypothesise that Sxl serves as a selective regulator of Pol III activity in neurons with high metabolic demand, such as mushroom body neurons (Placais *et al*., 2017; Mann *et al*., 2023). Future investigations using electrophysiology will be crucial to uncover the physiological role of *Sxl^RAC^*.

### Sxl^RAC^ moderates neuronal homeostasis

The Sxl^RAC^ isoform, an almost full-length protein expressed in male brains (Bopp *et al*., 1991; Cline *et al*., 2010; Moschall *et al*., 2018), carries a truncation in its N-terminus, which abolishes its splicing activity (Deshpande *et al*., 1999). While we observe a mild survival deficit in *Sxl* knockdowns, overexpression of *Sxl^RAC^* in adults results in a marked decline in survival (Fig. 5) and significant transcriptional changes associated with metabolic and translational processes (Fig. S4). These data underscore the importance of tightly regulating *Sxl* expression to maintain neuronal homeostasis. The contrasting effects of knockdown and overexpression (both at the phenotypic and transcriptomic levels) point to a dosage-sensitive role for Sxl in adult neurons, with context-dependent requirements. The lifespan alterations observed in overexpression models are likely explained as consequences of dysregulated tRNA synthesis and/or metabolic trade-offs (Beardmore *et al*., 2011; Niehues *et al*., 2016).

Using Small RNAseq and qPCR methods, it was discovered that *Sxl^RAC^*overexpression leads to changes in Pol III targets. Small RNAseq data indicates a bias toward upregulated tRNA biogenesis, particularly of lysine tRNAs (Fig. 5), along with effects in other small RNAs (Fig. 5). These findings align with recent discoveries in mammalian neurons, that highlight cell-type-specific tRNA expression in the brain and link it to the regulation of neuronal homeostasis (Kapur *et al*., 2024). Sxl^RAC^ isoform expression is likely dependent on the requirements of each cell, with downstream implications for translation and metabolism. This is further supported by OPP staining results, which reveal an overlap between cells expressing Sxl^RAC^ and those with increased translation, linking Sxl^RAC^ activity to local translational control in the brain. We therefore propose a model in which Sxl is recruited to chromatin to regulate Pol III activity in the adult male brain, regulating tRNA expression and boosting metabolism in high-activity cells. Given the emerging evidence that Pol III inhibition can improve ageing (Malik *et al*., 2024; Ureña *et al*., 2024), Sxl presents as a novel factor in age-related pathways.

### Sxl’s ancestral role

Our findings also raise intriguing questions about the conserved roles of Sxl and its orthologues across species. Though sexual dimorphism in *Sxl* expression is restricted to the *Drosophilidae* family, it has orthologues in other insect species, implying it may have evolved from a broader, sex-independent role, as seen in the Sxl^RAC^ isoform (Cline *et al*., 2010; Siera and Cline, 2008; Traut *et al*., 2006). The evolution of Sxl as a sex-specific gene (∼10 MY) likely reflects a later adaptation to regulate sexual differentiation (Cline *et al*., 2010). However, an ancestral, sex-independent function, such as regulating metabolic processes and RNA Pol III activity, may still be conserved. This raises the possibility that Sxl orthologues in other species, as well as in vertebrates, may regulate similar pathways, particularly in neuronal homeostasis. For instance, the ELAV family of RNA-binding proteins, which share sequence similarities with Sxl, also regulate RNA stability and translation in response to metabolic and stress signals (Lisbin *et al*., 2000; Pascale *et al*., 2005; Ince-Dunn *et al*., 2012; Kraushar *et al*., 2014), suggesting that similar regulatory mechanisms may be at play.

In conclusion, our study reveals a novel, non-splicing role for Sxl in regulating chromatin binding and Pol III transcription, with important implications for neuronal function, metabolism, and ageing. These findings provide new insights into the broader functional repertoire of Sxl and underscore the potential to explore its conserved roles across species, from insects to vertebrates. Further research into the mechanisms governing Sxl’s chromatin recruitment and its interaction with Polr3E will be crucial for understanding its full range, particularly in the context of neuronal function.

## Materials and Methods

### Drosophila husbandry

Crosses were established and maintained under conditions optimised for each experiment. For larval experiments, *elavC155-GAL4* virgin females were crossed to Dam lines (see below) to drive expression in early-developing neurons. For adult-specific manipulations, *w^-^; tub-GAL80^ts^; nSyb-GAL4* virgin females were crossed to RNAi lines and corresponding controls: *UAS-Sxl-RNAi-I* (VDRC #109221), *UAS-Sxl- RNAi-II* (BDSC #34393), *UAS-Polr3E-RNAi* (BDSC #55656), *y; RNAi-TK* (VDRC #60101), *y; p(UAS-CaryP)attP2* (BDSC #36303), *y;p(UAS-CaryP)attP40* (BDSC #36304). Crosses were maintained at 18_°C throughout development, with offspring shifted to 29_°C to induce GAL4-mediated expression. The *Sxl-T2A-GAL4/FM7* line (kindly provided by Jan Madenbach) was used to visualize *Sxl* expression in the adult brain. For localisation and colocalisation experiments, these flies were crossed to *10xUAS-IVS-mCD8::GFP* (BDSC #32185) and *y; p(UAS-mCherry)attP2* (BDSC #35787) males, respectively.

### *Drosophila* transgenics

For generating *UAS*-Dam-splice factor constructs, *Sxl, U1, tra, U2AF50* and *X16* coding sequences were amplified by PCR from cDNA, digested with NotI and XbaI and Gibson assembly cloned (using NEBuilder HiFi DNA Assembly kit (NEB)) into *pUASTattB-LT3-Dam* (Southall *et al*., 2013). For *Sxl^RNA^*, 3 overlapping segments of *Sxl* were amplified with mismatch primers (to mutate the RNA binding sites) and simultaneously Gibson cloned into *pUASTattB-LT3-Dam. Sxl^GS^* was generated by gene synthesis (Integrated DNA Technologies (IDT)) and cloned into *pUASTattB- LT3-Dam*. *For generating UAS-Sxl^RAC^*, the coding sequence was PCR amplified from cDNA and Gibson cloned into *pUASTattB*. To generate transgenics, these constructs were injected into *attP2* flies (*y w P{y[+t7.7]=nos-phiC31\int.NLS}X #12;; P{y[+t7.7]=CaryP}attP2 –* Department of Genetics Microinjection Service, Cambridge) for integration on the third chromosome. In functional studies, experimental flies were compared to flies landing-site control flies *y; p(UAS-mCherry)attP2* (BDSC #35787) crossed with the same driver.

### Targeted DamID

Dam fusion and Dam-only males were crossed to *elav^C155^*-GAL4 virgins for expression of Dam in neurons. 50 larval central nervous systems were dissected and stored at -80°C. Genomic DNA extraction, methylated DNA enrichment and library preparation were carried out as previously described (Marshall *et al*., 2016). Two biological replicates were generated for each experiment. For the experiment profiling Sxl binding in neurons with *Polr3E* RNAi (BDSC #55656), the larvae were raised at 18°C to prevent premature lethality. 3 replicates were used for each condition (for *wild type*, x1 female only sample and x2 mixed sex samples, and for *Polr3E* RNAi, x3 mixed sex samples – the female only sample is comparable to the mixed sex samples – see correlation coefficients in Table S8).

Libraries were sequenced with 50 bp single end sequencing on Illumina HiSeq platform. >10 million reads were obtained for each library. Data were processed using a previously described Perl pipeline (Marshall and Brand, 2015) (https://github.com/owenjm/damidseq_pipeline).

Peak calling was performed using a previously described Perl program (available at https://github.com/tonysouthall/Peak_calling_DamID) which allows for the identification of broadly bound regions that characterise DamID data (Estacio-Gomez *et al*., 2020). Briefly, false discovery rate (FDR) was calculated for regions consisting of >1 GATC bounded fragments for each replicate. Significant (FDR<0.01%) regions present in all replicates are merged to form a final peak file.

### Immunohistochemistry

Immunohistochemistry was performed on third instar larval and adult central nervous systems (CNS). Dissections were carried out in 1× PBS, followed by fixation in 4% formaldehyde (methanol-free; Polysciences Inc.) diluted in PBS for 20 minutes at room temperature. Tissues were washed three times in PBST (0.3% Triton X-100 in PBS) for 5 minutes each, then blocked in 10% normal goat serum (NGS) in PBST for 1 hour at room temperature. Samples were incubated with primary antibodies overnight at 4_°C. The following day, tissues were washed three times in PBST and incubated with secondary antibodies diluted in PBST for 2 hours at room temperature, followed by final washes. Samples were mounted in VECTASHIELD Antifade Mounting Medium (Vector Laboratories) on standard glass slides. Primary antibodies used were mouse anti-Repo (1:500; 8D12 Developmental Studies Hybridoma Bank, DSHB) and rat anti-Elav (1:500; 7E8A10 DSHB). Secondary antibodies were Alexa Fluor 545 and 633 (1:200; Thermo Fisher Scientific). Images were subsequently analysed using a Zeiss LSM 510 confocal microscope.

### Behavioural and phenotypic assays

Survival assays were conducted as previously described (Hassan *et al*., 2020). Progeny from appropriate genetic crosses were reared at standardized larval densities. Upon eclosion, adult flies were allowed to mate for 24_h before being separated by sex. For each experimental condition, groups of 10 flies of the relevant genotype were transferred into vials containing fresh food. Vials were replaced every two days, and mortality was recorded at each transfer. To circumvent developmental effects, flies were reared at 18_°C and shifted to 29_°C post-eclosion to induce GAL4-driven expression by inactivating GAL80^ts^. Each condition was assessed with a minimum of 10 biological replicates to generate survival curves, which were compared to appropriate controls. Statistical analyses were performed using GraphPad Prism (v5.0). Significance was determined using the Gehan-Breslow-Wilcoxon and Log-rank (Mantel Cox) tests.

Age-related changes in locomotor performance were assessed using the Reactive Iterative Negative Geotaxis (RING) assay. A custom-designed apparatus was employed to ensure consistent starting conditions, with all flies dropped from a uniform height of 15 cm at a controlled velocity. For each experimental condition, three biological replicates were assessed, each consisting of 10 flies placed in an empty plastic vial. Flies were allowed to acclimate for 5 minutes prior to testing. Each group underwent three technical replicate trials, with 2-minute rest intervals between drops. Climbing behaviour was recorded through video using a handheld camera.

Performance was quantified by calculating the percentage of flies reaching the halfway mark (4 cm) within 10 seconds. Data were analysed using two-way ANOVA, followed by appropriate post hoc tests.

### RNA-seq and data analysis

For each experimental condition, three biological replicates were prepared, with total RNA extracted from the heads of 10-20 flies per replicate using TRIzol reagent (Thermo Fisher Scientific), according to the manufacturer’s instructions. RNA sequencing and small RNA library preparation and sequencing were performed by Genewiz (Azenta Life Sciences), generating ∼20 million paired-end reads per sample.

Raw RNA-seq data were processed using the Galaxy platform (https://usegalaxy.eu). Adapter sequences were trimmed from FASTQ files using Trimmomatic (v0.39) with the ILLUMINACLIP parameter, and read quality was assessed using FastQC (v0.74). Reads were aligned to the *Drosophila* reference genome (release r6.58) using HISAT2 (v2.2.1). Gene-level quantification was performed with featureCounts (v2.0.8), and differential gene expression was analysed using DESeq2 (v2.11.40.8). Gene Ontology (GO) enrichment analysis was conducted using tools provided by the Gene Ontology Consortium and PANTHER classification system.

For small RNA-seq, data were processed independently using the Galaxy platform. Adapter trimming was performed with Trim Galore! (v0.6.7), and read quality was assessed using Falco (v1.2.4). Only Read 1 (R1) was used, as small RNAs are typically captured in R1, and Read 2 (R2) was excluded due to persistent adapter contamination errors in processing after trimming. Reads were mapped to the *Drosophila* reference genome (r6.58) using Bowtie2 (v2.5.3), and small RNA species, including tRNAs, were quantified using featureCounts (v2.0.8). Differential expression analysis was again performed using DESeq2 (v2.11.40.8), and data visualisation, including volcano plots, was completed in R (v4.2.2) using the ggplot2 package.

### Quantification of RNA Polymerase III targets

Following quantitative RNA extraction, reverse transcription was performed using the iScript cDNA Synthesis Kit (Bio-Rad) with random hexamers, in accordance with the manufacturer’s instructions. Quantitative real-time PCR (qRT-PCR) was conducted using iTaq Universal SYBR Green Supermix (Bio-Rad). Expression of target genes was normalised to U3, a small nucleolar RNA transcribed by RNA polymerase II and compared to expression levels in the corresponding control samples. Primers targeting pre-tRNA species were based on previously validated sequences (Frendewey *et al*., 1985; Chan *et al*., 2016; Malik *et al*., 2024); all primer sequences are listed below.

pre-tRNAHisGTG-1 F CGTGATCGTCTAGTGGTTAG, pre-tRNAHisGTG-1 R CCCAACTCCGTGACAATG, pre-tRNAIleTAT-1 F CGCACGGTACTTATAATCAG, pre-tRNAIleTAT-1 R, CCAGGTGAGGCTCGAACTC, pre-tRNALeuCAC-1 F GCGCCAGACTCAAGATTG, pre-tRNALeuCAC-1 R TGTCAGAAGTGGGATTCG, U3 forward CACACTAGCTGAAAGCCAAG, and U3 reverse CGAAGCCCTGCGTCCCGAAC.

For each genotype, sample sizes were too small to reliably assess normality; therefore, a one/two-tailed Mann–Whitney test was used to account for the non- parametric nature of the data.

### O-Propargyl-Puromycin (OPP) assay

Protein synthesis was assessed using the Click-iT Plus OPP Assay Kit (Thermo Fisher Scientific), following the manufacturer’s instructions. Dissections were performed in Schneider’s Insect Medium to preserve neural tissue integrity. Isolated brains were incubated with O-propargyl-puromycin (OPP) for 1 hour at room temperature to label nascent peptides. Following incubation, samples were fixed in 4% paraformaldehyde (PFA) for 15 minutes, then permeabilised and washed in 0.5% Triton X-100 in PBS (PBST). Click chemistry-based detection was carried out using a reaction cocktail composed of 880 µL 1× reaction buffer, 20 µL copper protectant, 100 µL 1× additive, and 2.5 µL Alexa Fluor picolyl azide. After the labelling reaction, samples were washed in 0.5% PBST. Samples were mounted in Vectashield mounting medium and imaged using a Leica Stellaris 5 light sheet microscope (LS1).

Colocalisation analysis was performed using ImageJ (v1.54p) with the Colocalisation Threshold plugin. Images were imported as 16-bit files, split into individual channels, and processed using the *Enhance Background* function across all slices with stack histogram enabled. No regions of interest (ROIs) were selected to avoid location bias, and Costes’ automatic thresholding was applied to eliminate background signal. Overall *R*coloc value was calculated by averaging across eight brains.

### Data availability

All raw sequence files and processed files have been deposited in the National Center for Biotechnology Information Gene Expression Omnibus (GSE294838). This is a SuperSeries containing GSE239790 (Targeted DamID), GSE294835 (RNA-seq) and GSE294837 (small RNA-seq).

## Supporting information

All supplementary files

## Acknowledgements

Thanks to Jan Medenbach for providing the *Sxl-T2A-GAL4* transgenic line, Nazif Alic for advice on qPCRs targeting pre-tRNAs, and Julia Bandiak for initial explorations and stock building. Furthermore, thank you to the entire Southall lab for helpful discussions and pre-reading the manuscript. We also thank the Bloomington Drosophila Stock Center (NIH P40OD018537) and the Vienna Drosophila Resource Center (VDRC, www.vdrc.at) for fly stocks. This work was funded by a Wellcome Trust Investigator grant 104567/Z/14/Z to T.D.S. and a BBSRC grant BB/X00256X/1 to T.D.S. and J.v.d.A.

## Supplementary Figures

**Figure S1.**
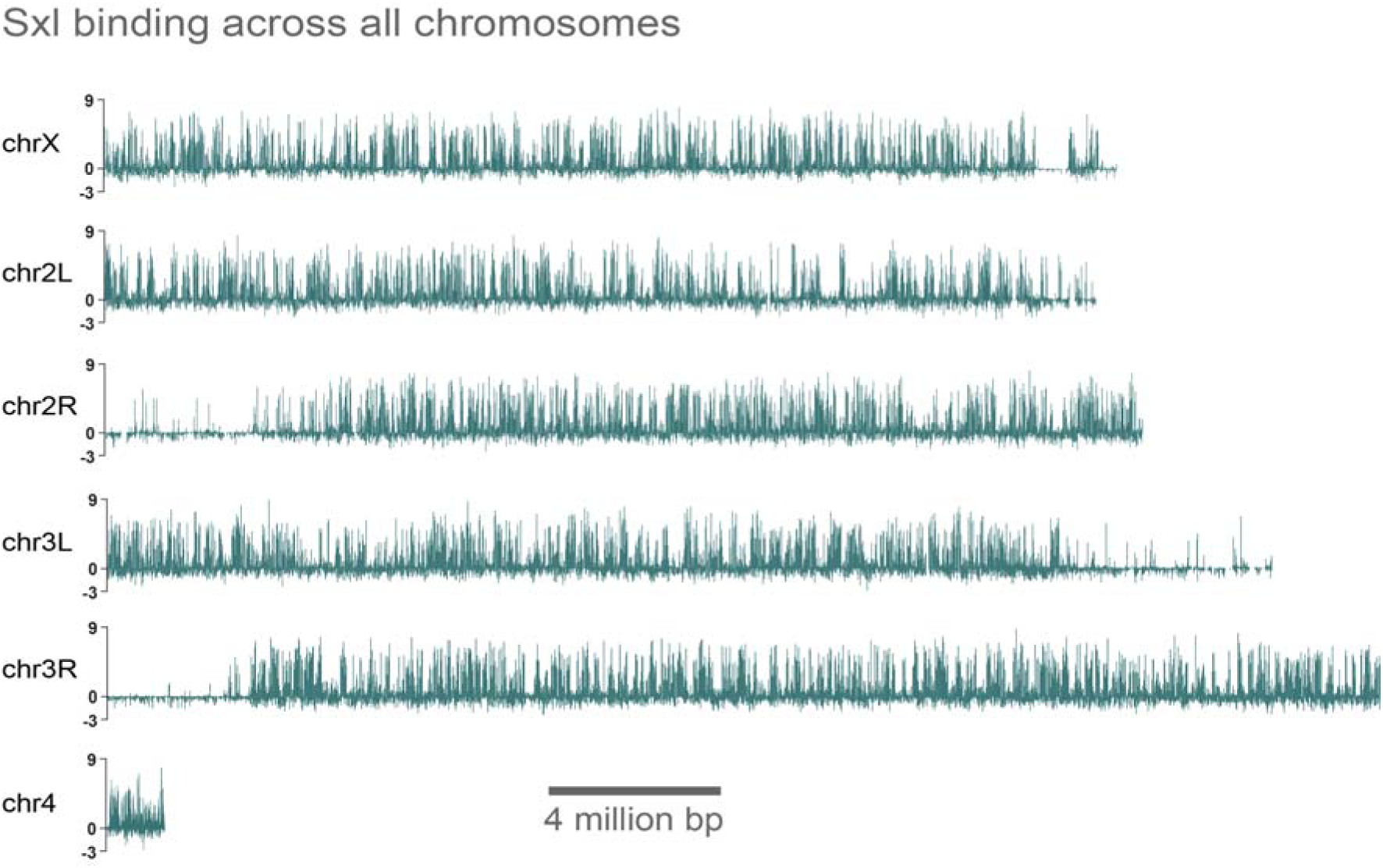
Sxl robustly binds chromatin across the genome. TaDa profiles for *Sxl* across chromosomes X, 2 L/R, 3 L/R and 4. The *y*-axis represents the log_ ratio of Dam–Sxl binding over that of Dam-alone.

**Figure S2.**
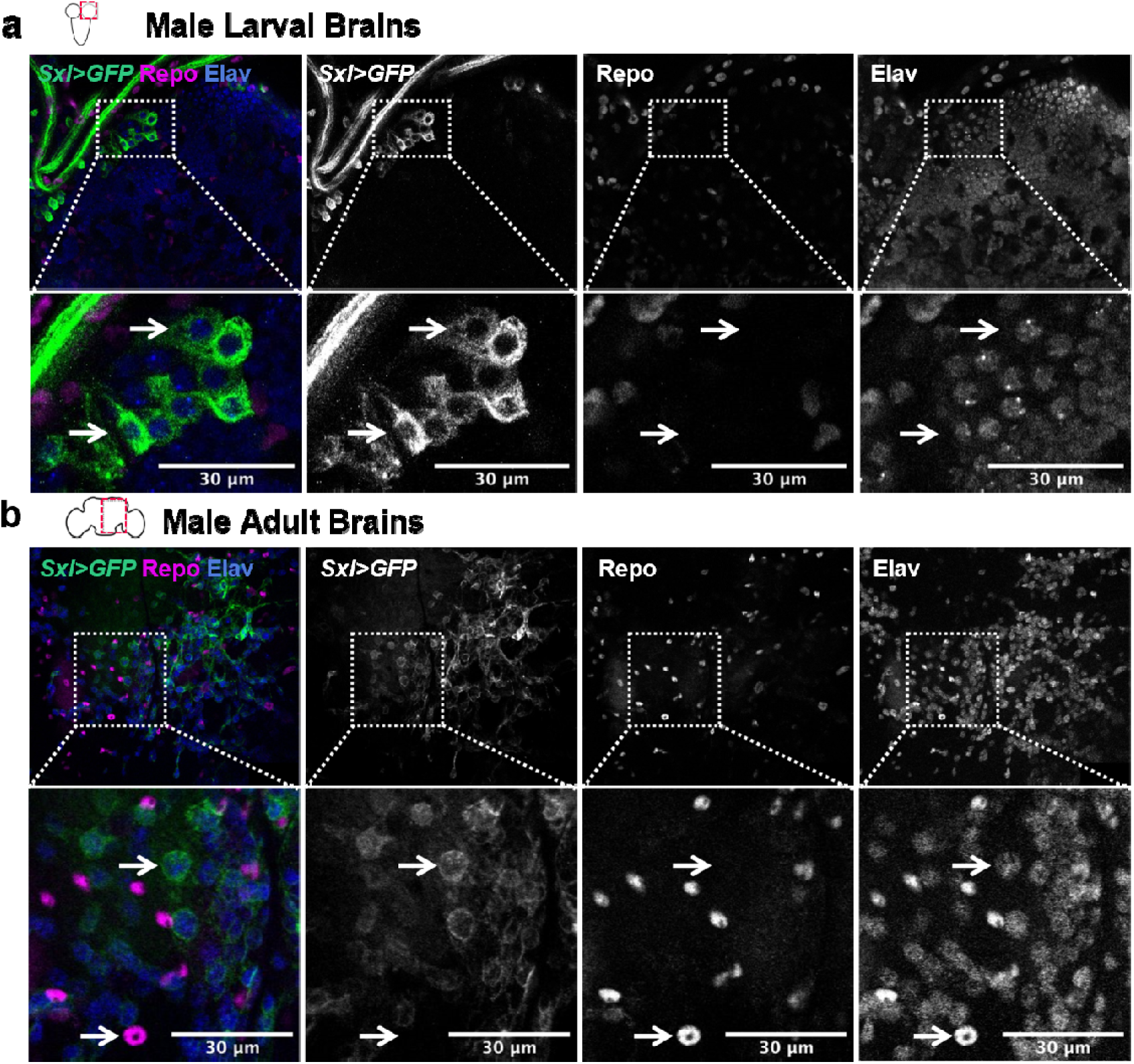
Sxl is expressed in the neurons of the male brain. **a**, Immunolabelled sections of the larval male brain showing *Sxl* expression (mcd8-GFP, green), glial marker repo (magenta) and neurons Elav (blue)**. b**, Equivalent labelling in adult brain sections. White arrows indicate *Sxl*-positive cells co-localised with Elav but not repo, consistent with neuron-specific expression. Images captured at 40x magnification; scale bar, 30μm.

**Figure S3.**
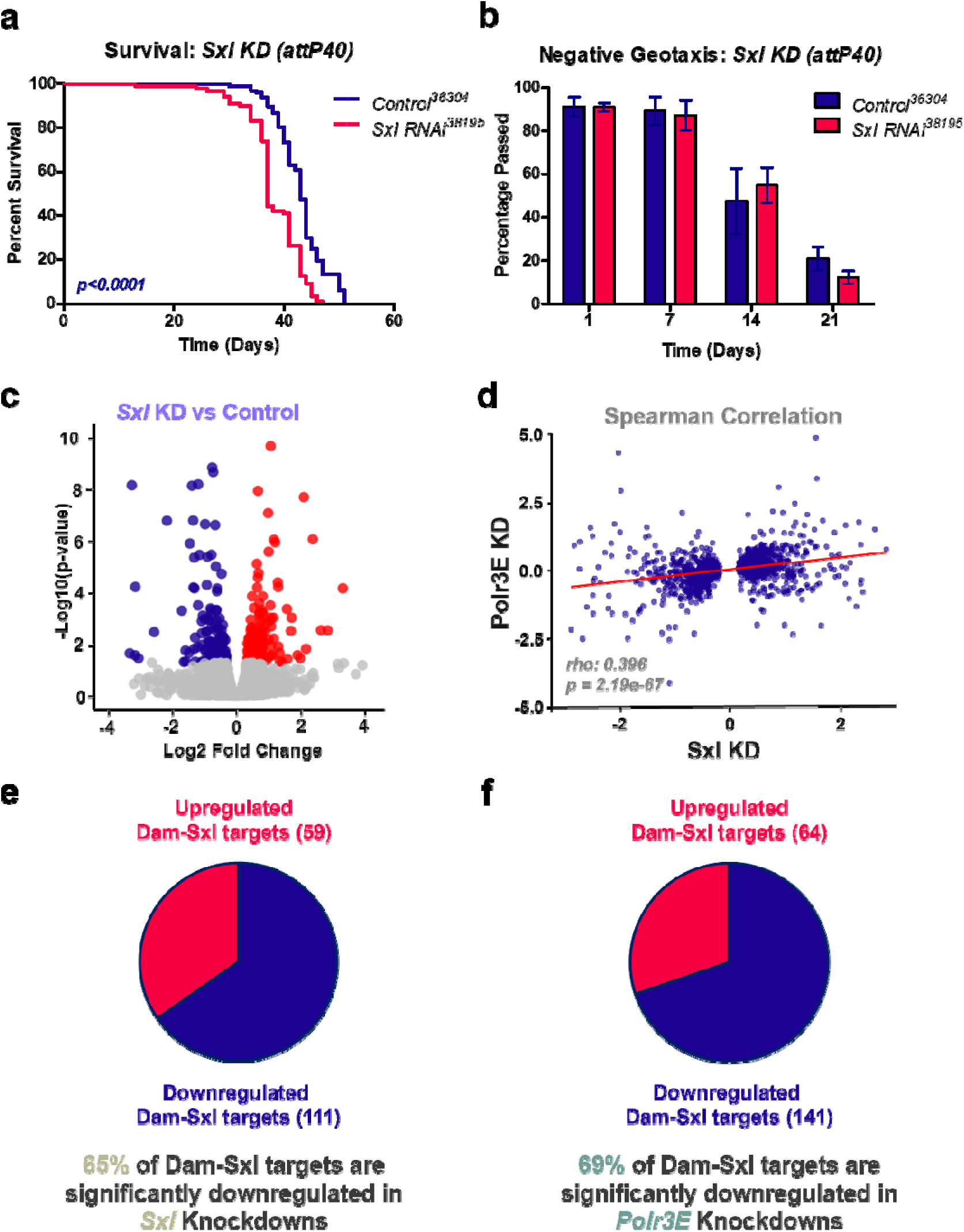
Sxl knockdown phenocopies Polr3E Loss in adult males. **a**, Survival curves of adult males following pan-neuronal knockdown of *Sxl* using a n independent RNAi line (BDSC #38195) compared to RNAi control (BDSC #36304) (n>90 per group). Statistical significance assessed using Gehan–Breslow–Wilcoxon and Kaplan-Meier test. **b**, Negative geotaxis assay showing no significant changes in climbing ability following *Sxl* knockdown using the same RNAi line. The *y*-axis indicates the percentage of flies surpassing the 4cm midpoint. **c**, Volcano plot displaying differential gene expression after *Sxl* knockdown in adult male neurons (VDRC #109221) relative to controls (VDRC #60101). Significantly upregulated transcripts (*p<*0.05) are shown in red; downregulated transcripts (*p<*0.05) in blue. **d**, Spearman correlation analysis of transcriptional changes in *Sxl-* and *Polr3E-*depleted neurons reveals a significant positive relationship (ρ = 0.396, *p*<0.0001). **e**, Pie chart illustrating that 65% of Dam-Sxl chromatin targets are downregulated upon *Sxl* knockdown in adult male neurons. **f**, Equivalent analysis indicating 69% of Dam-Sxl targets are similarly downregulated following *Polr3E* knockdown. Data underlying this figure are available in Tables S19, S20, S21, S22, S23, S24, S25, S26, S27 and S28.

**Figure S4.**
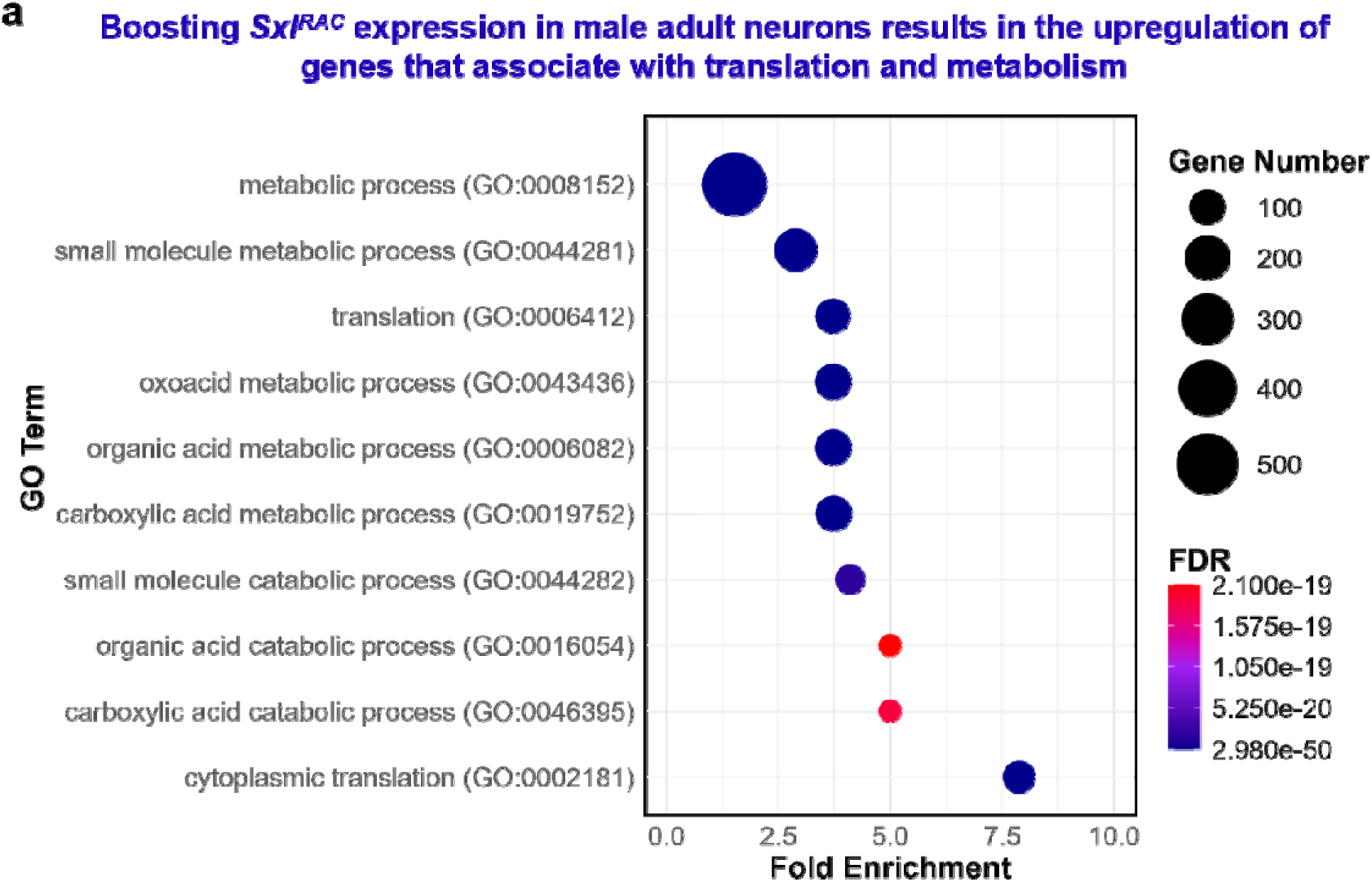
Elevated Sxl^RAC^ leads to upregulation of genes that associate with metabolism and translation. **a**, Gene ontology enrichment analysis of transcripts downregulated after boosted *Sxl^RAC^* expression in adult male neurons. Dot plot shows the top 10 GO terms ranked by fold enrichment (*x*-axis); dot size indicates the number of gene hits, and color reflects FDR. Data underlying this figure are available in Tables S29, S30 and S31.

**Figure S5.**
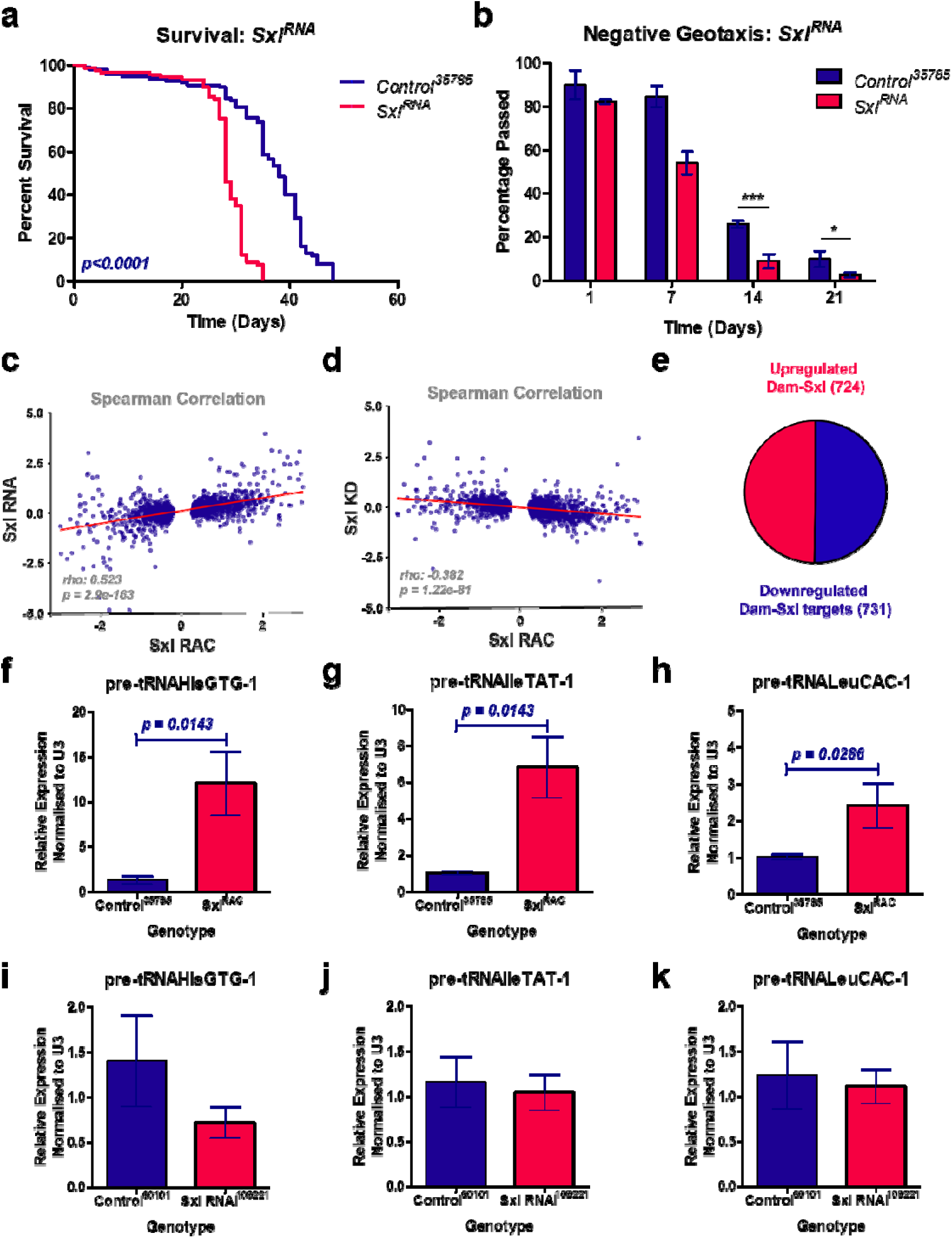
Elevated Sxl^RAC^ leads to altered tRNA synthesis. **a**, Survival curves of adult males following pan-neuronal expression of *Sxl^RNA^*, compared to *mCherry* controls (n > 99 per group). Statistical significance assessed using Gehan–Breslow–Wilcoxon test. **b**, Negative geotaxis assay reveals impaired climbing in males overexpressing *Sxl^RNA^*. The *y*-axis indicates the percentage of flies surpassing the 4cm midpoint. Data were analysed by two-way ANOVA with Bonferroni post hoc correction; \**p*<0.05 and \*\*\**p*<0.001. **c**, Spearman correlation analysis of transcriptional changes in *Sxl^RAC^*- and *Sxl^RNA^*-expressing neurons shows a significant positive relationship (ρ=0.523, *p=*2.9e-163). **d**, In contrast, a significant negative correlation is observed between *Sxl^RAC^* overexpression and *Sxl knockdown profiles* (ρ=-0.382, *p=*1.22e-81). **e**, Pie chart illustrating an even split of Dam-Sxl chromatin targets being up or downregulated in *Sxl^RAC^*-expressing males. **f-h**, Overexpression of *Sxl^RAC^* in male neurons increases pre-tRNA abundance. **f**, Relative expression of *pre-tRNA^His^* measured by qPCR (n=4, *p*=0.0143, Mann– Whitney test). **g**, Relative expression of *pre-tRNA^TAT^* (n=4, *p*=0.0143). **h**, Relative expression of *pre-tRNA^CAC^* (n=4, *p*=0.0286). **i-k**, Pre-tRNA levels trend lower in *Sxl* RNAi-expressing heads (VDRC #109221). **i**, Relative expression of *pre-tRNA^His^* (n=6, ns). **j**, Relative expression of *pre-tRNA^TAT^* (n=6, ns). **k**, Relative expression of *pre-tRNA^CAC^* (n=6, ns). Data underlying this figure are available in Tables S32, S33, S34, S35, S36, S37, S38, S39, S40, S41.

**Figure S6.**
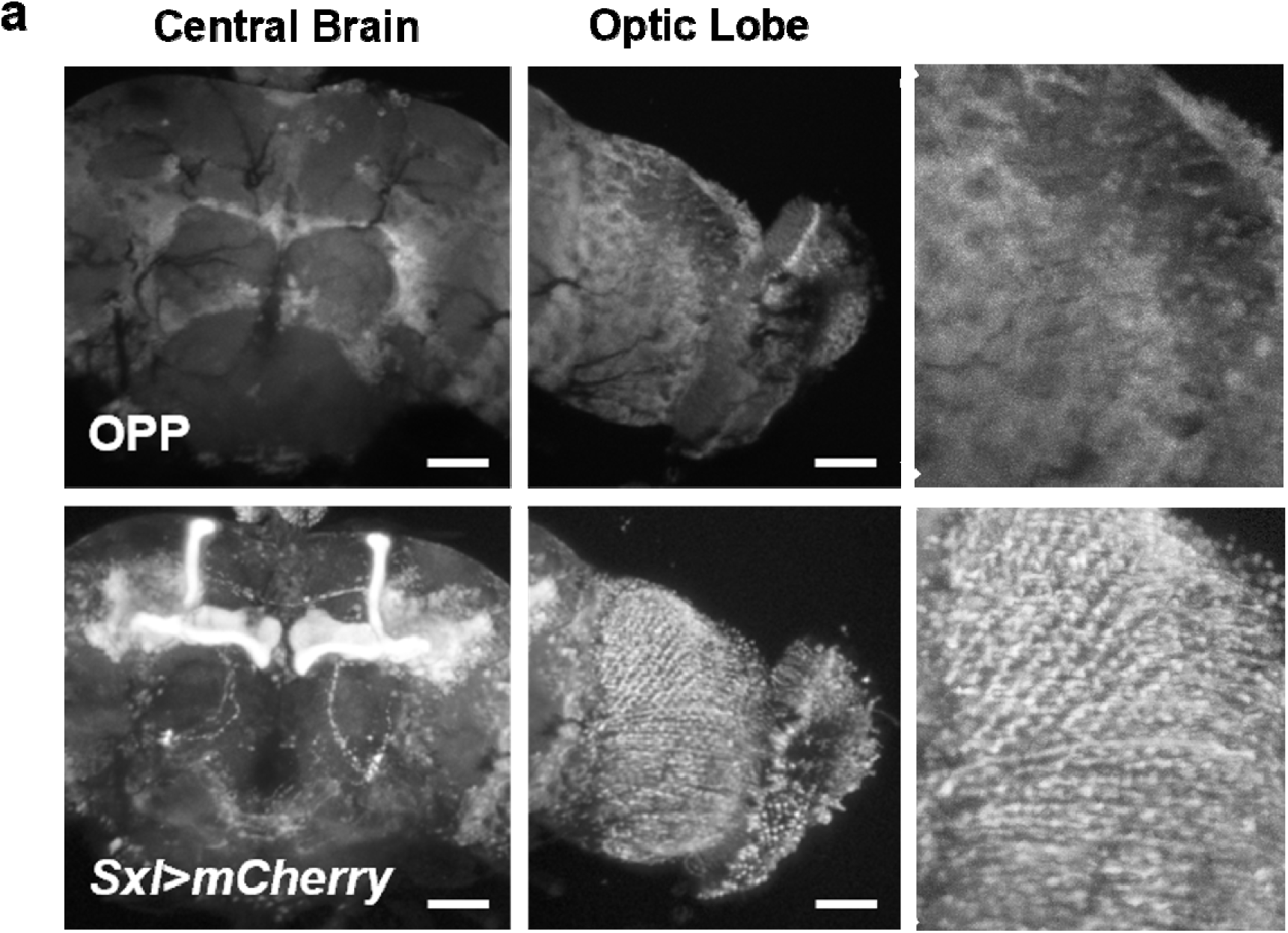
Neuronal regions with enriched Sxl expression exhibit elevated protein synthesis. **a**, Fixed adult brain section labelled with O-propargyl-puromycin (OPP) to visualise nascent protein synthesis, alongside mCherry expression driven by *Sxl-T2A-GAL4* (*Sxl>mCherry*). Regions of elevated protein synthesis closely align with areas of enriched Sxl expression, notably within the mushroom bodies and medulla. Images acquired at 20x magnification; scale bar, 30_μm.

