## Supplementary figures and images for "Sex-lethal is recruited to chromatin to promote neuronal tRNA synthesis in males through RNA Polymerase III regulation"

### S22_Sxl_RNAi_I_RNAseq_Plots.pdf

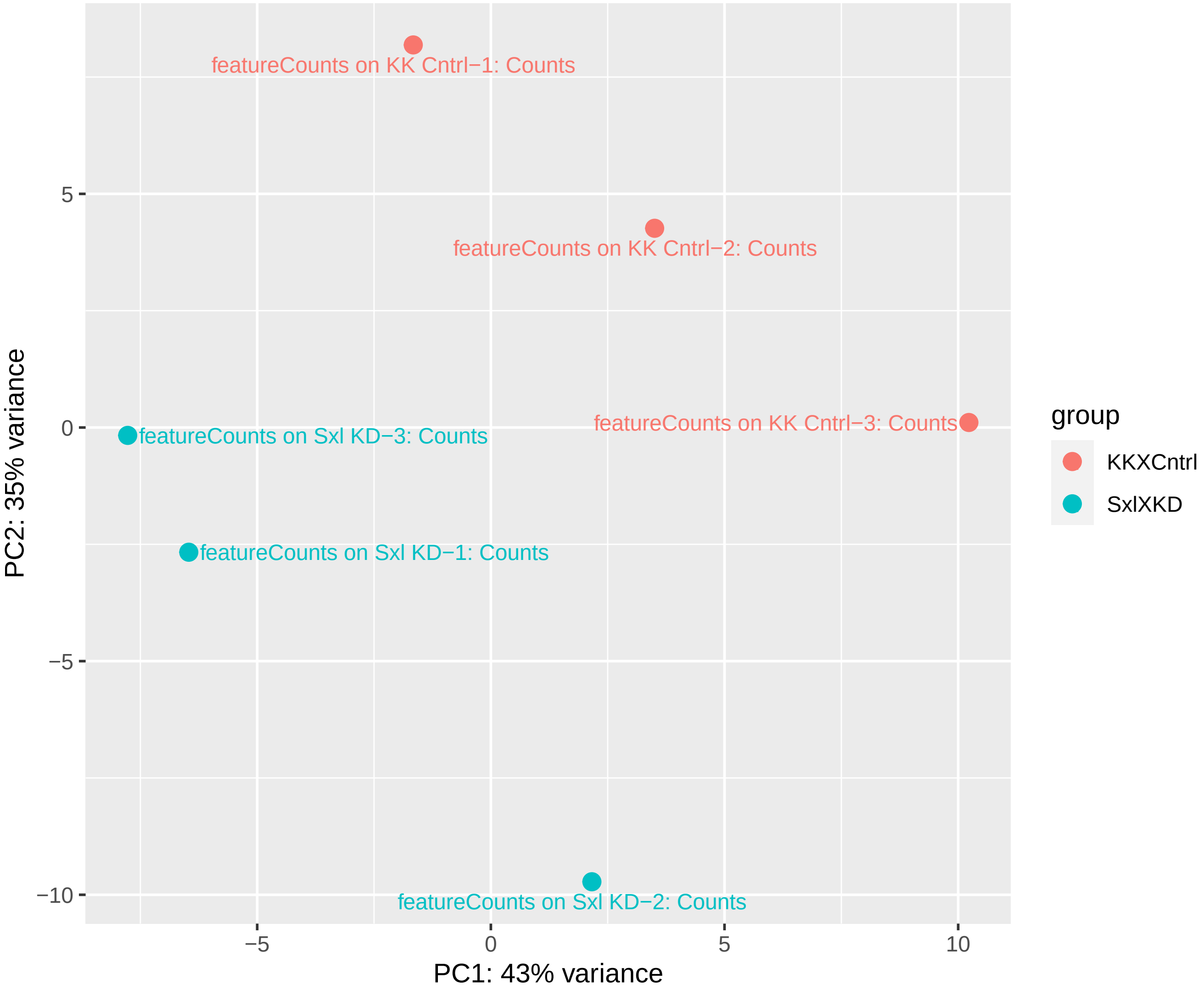

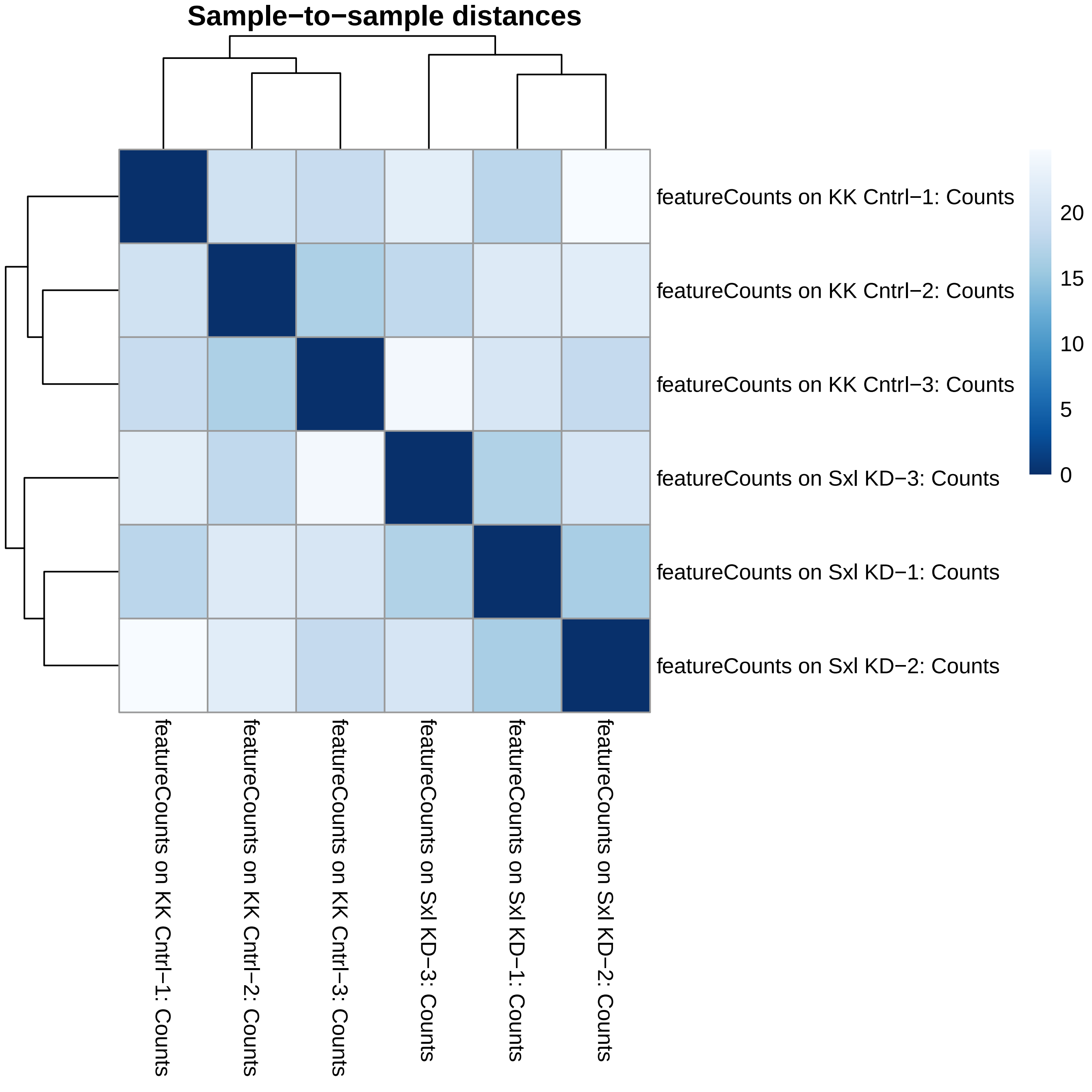

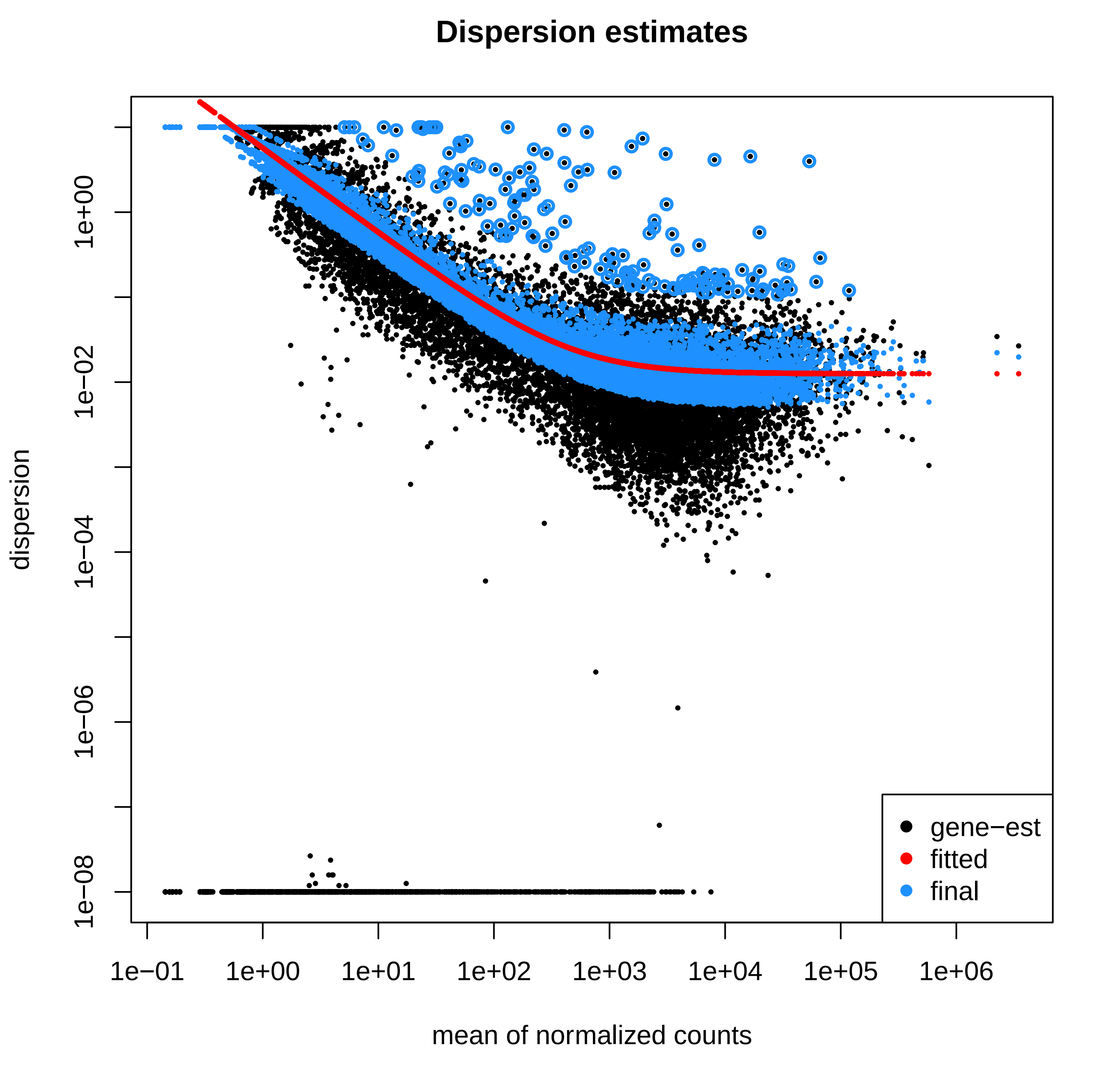

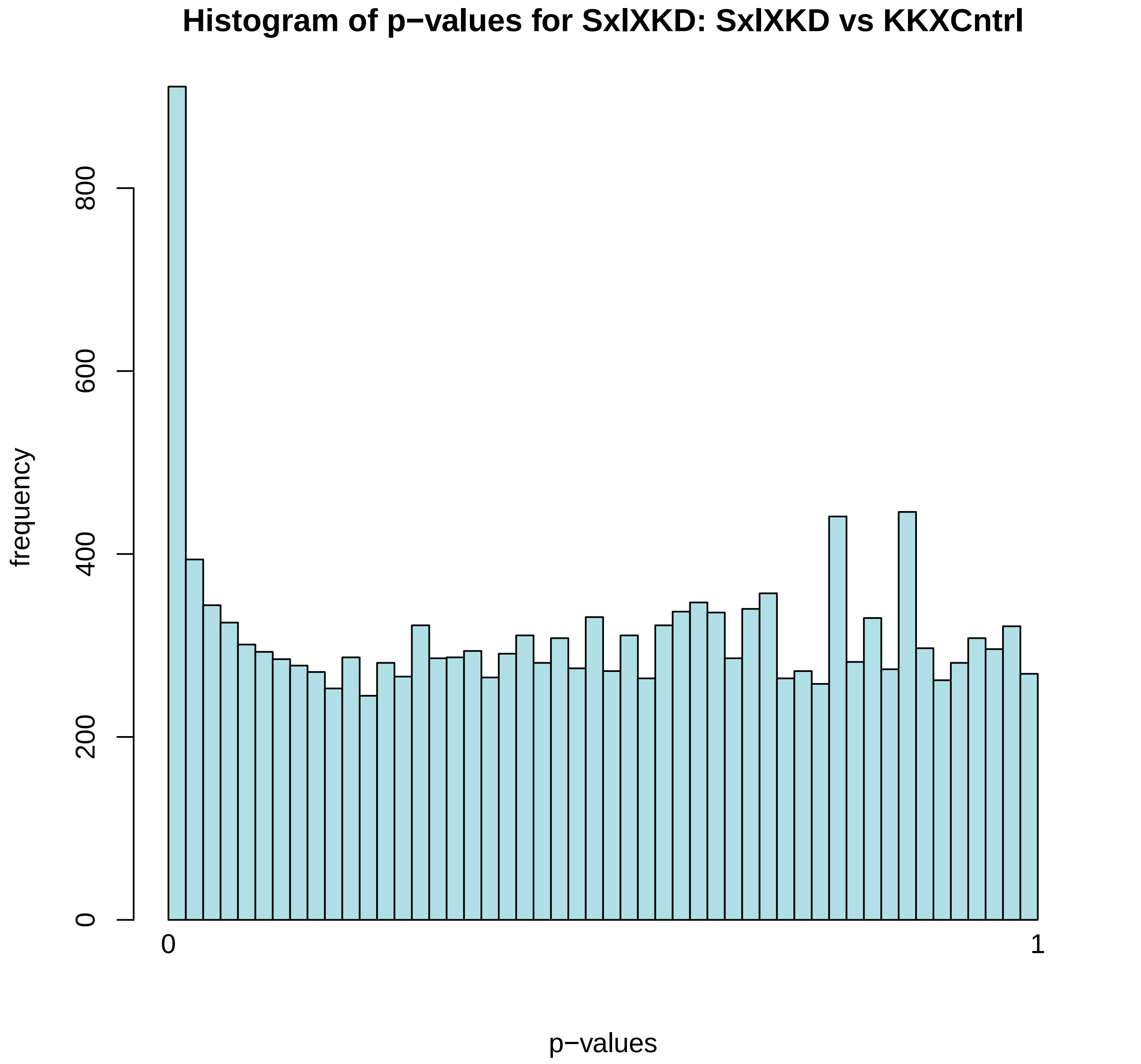

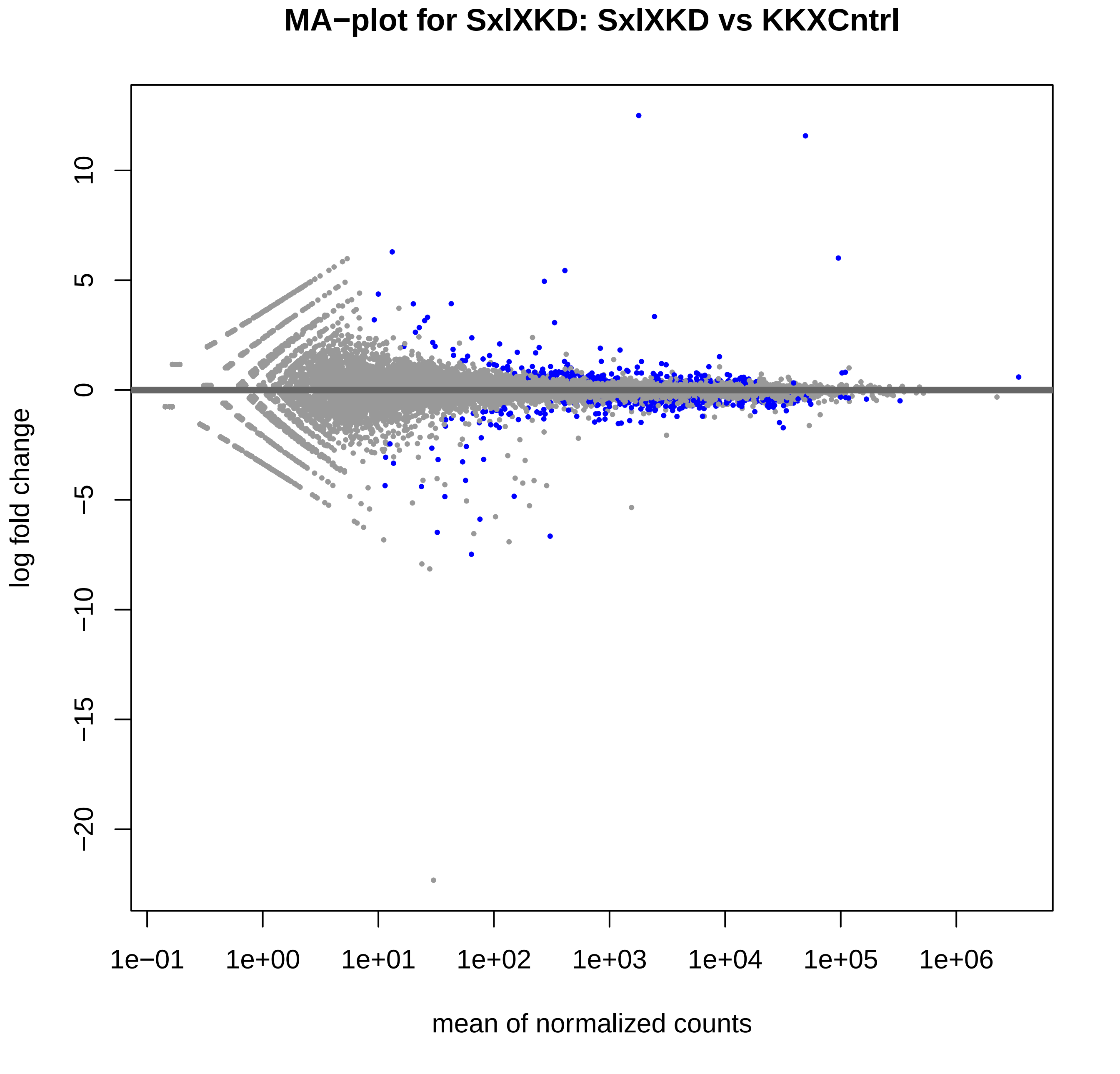

### S25_Polr3E_RNAi_I_RNAseq_Plots.pdf

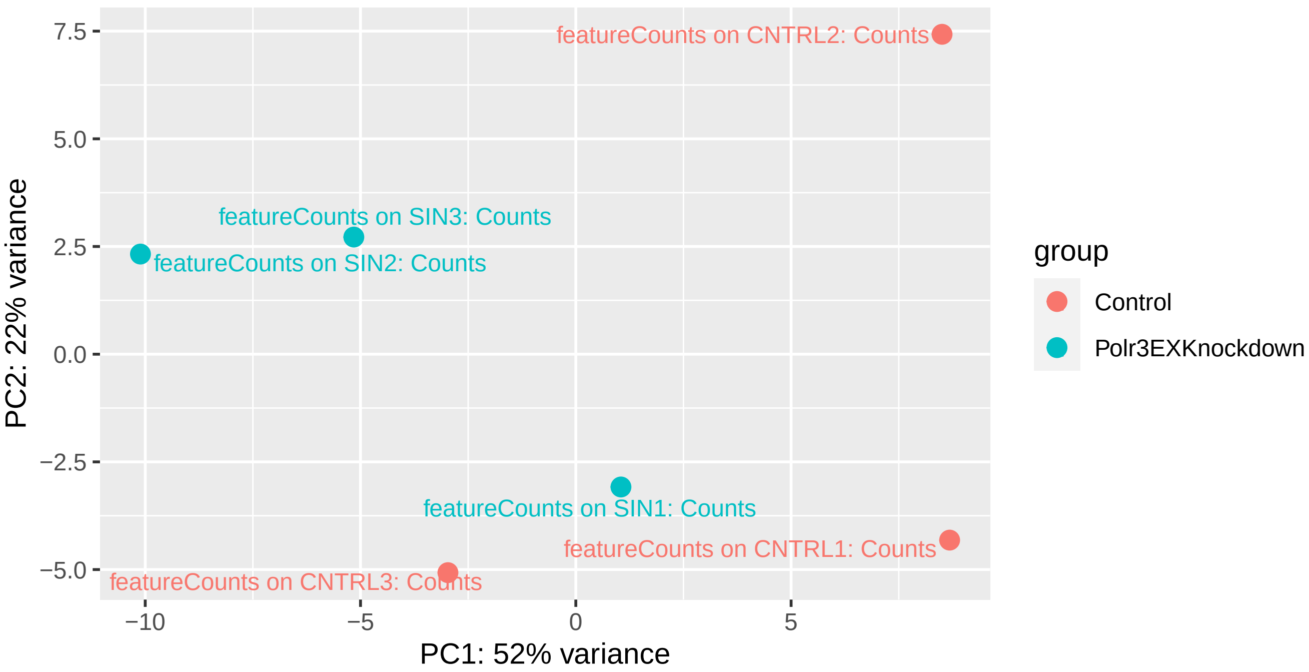

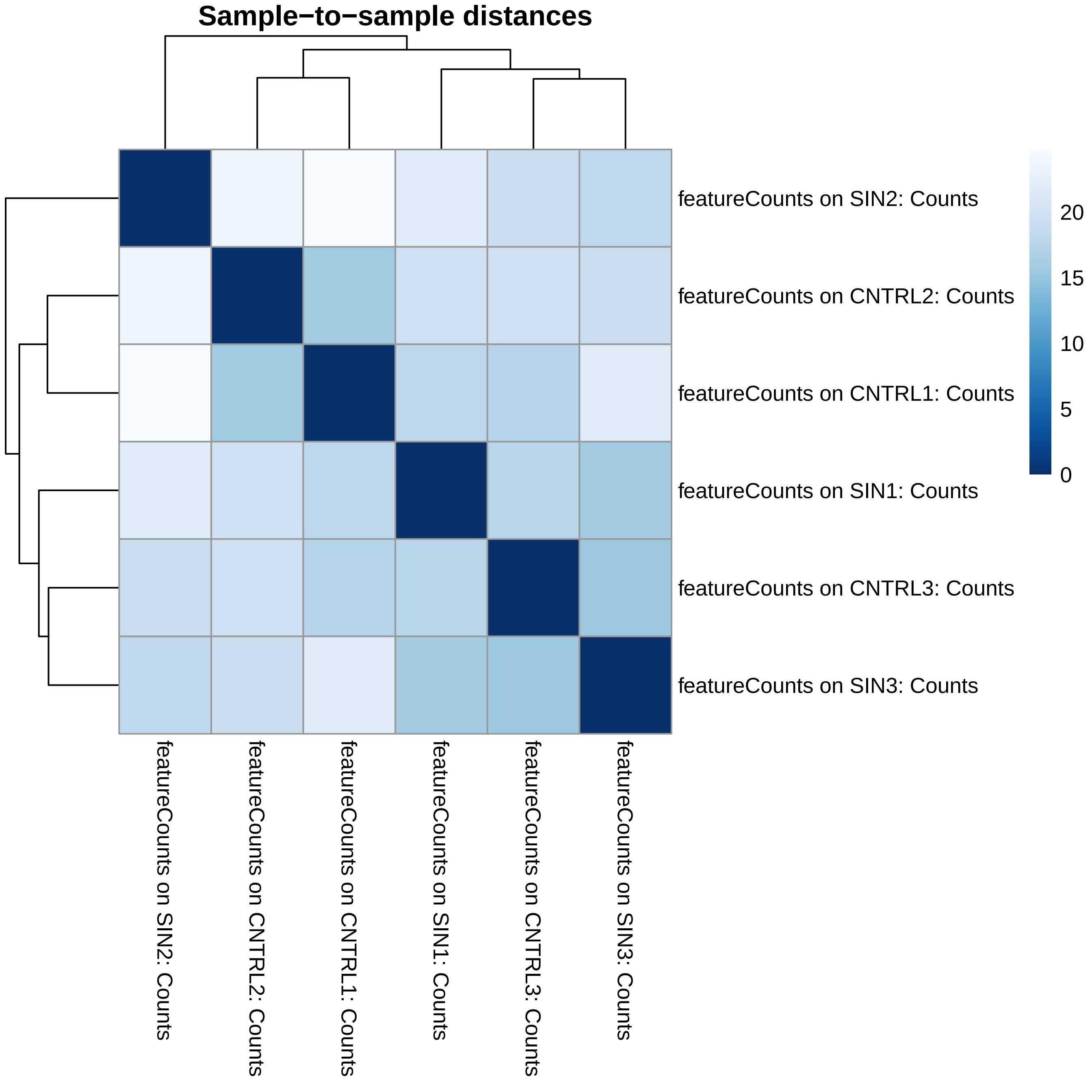

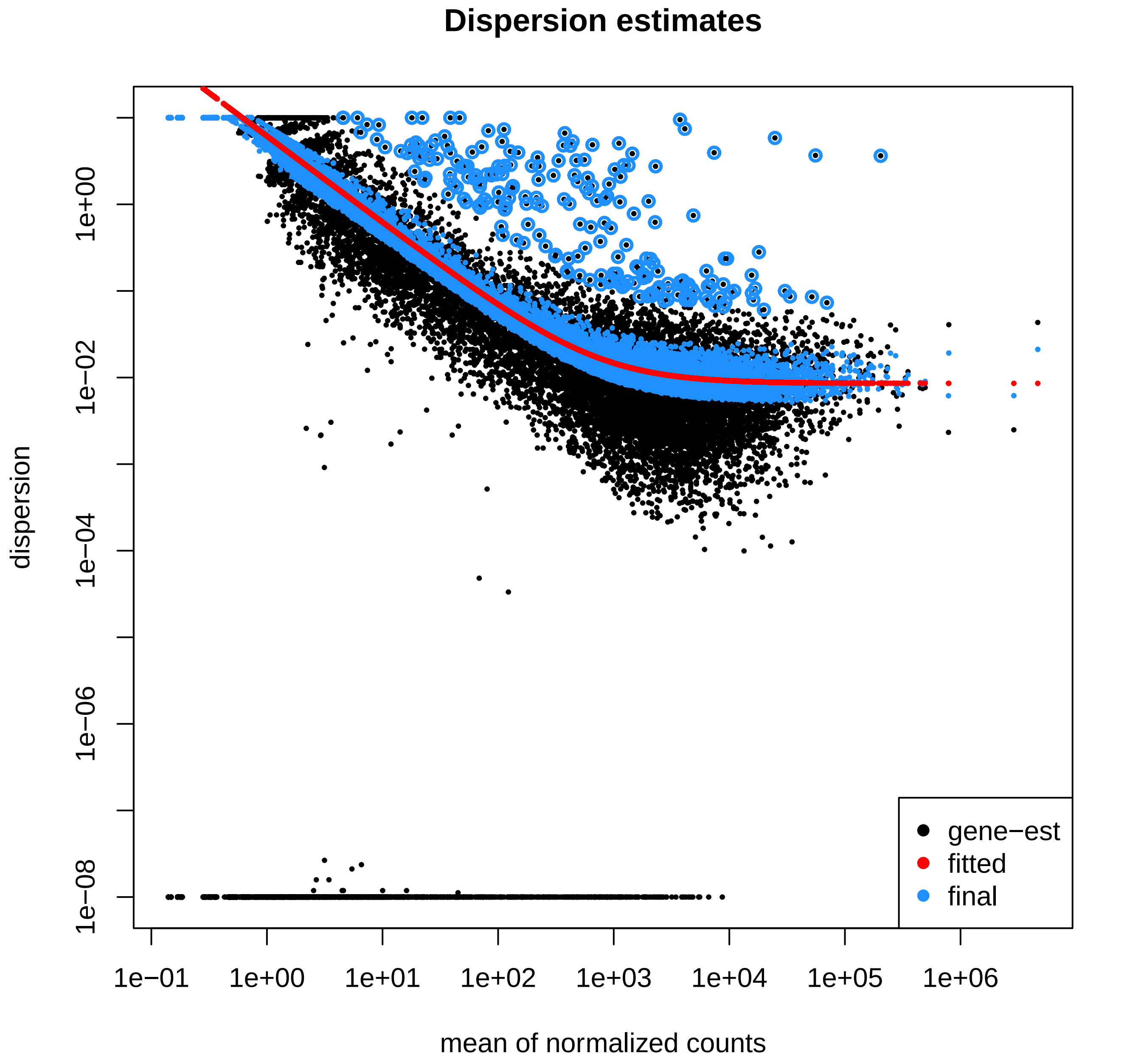

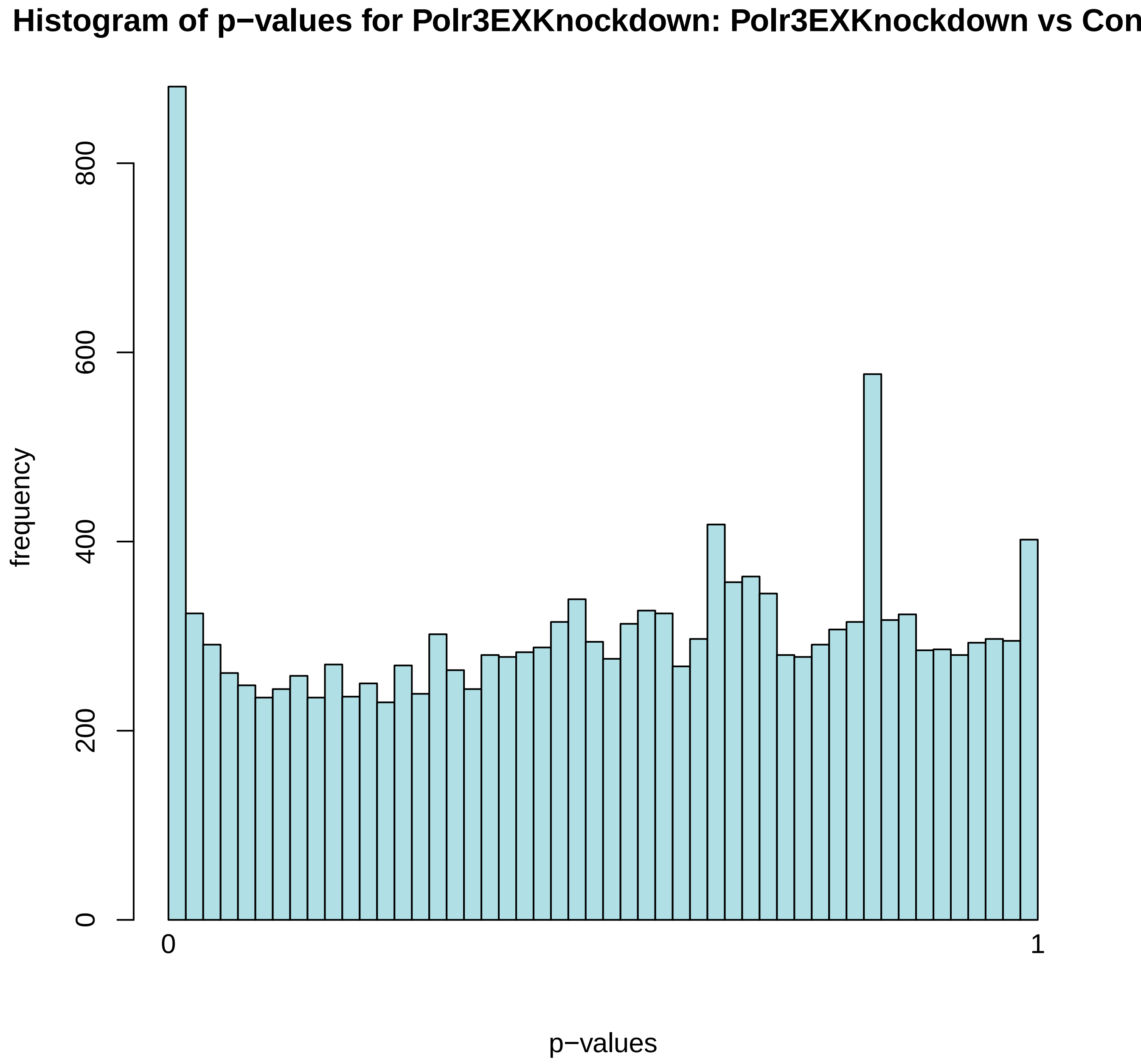

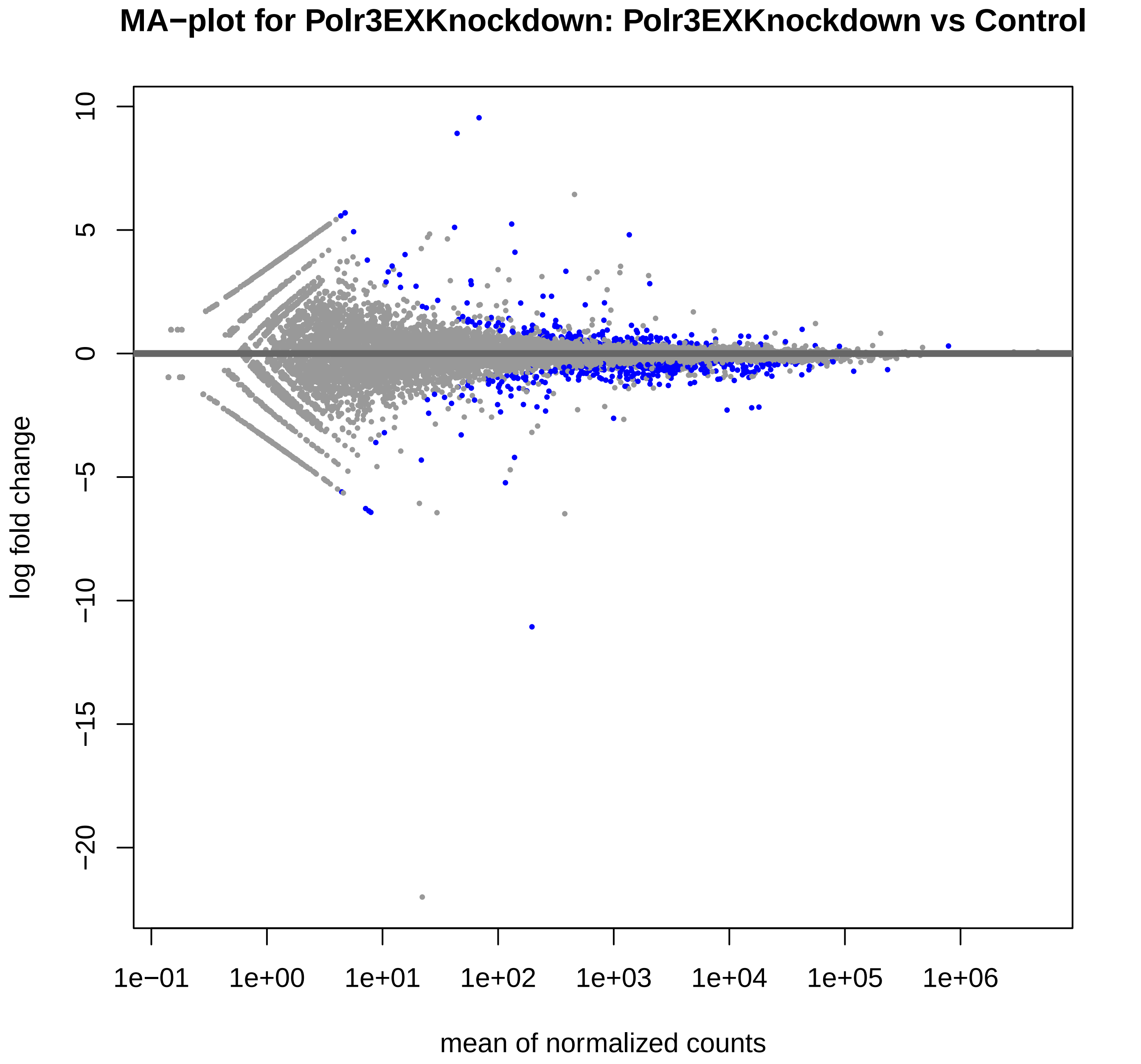

### S30_Sxl_RAC_RNAseq_Plots.pdf

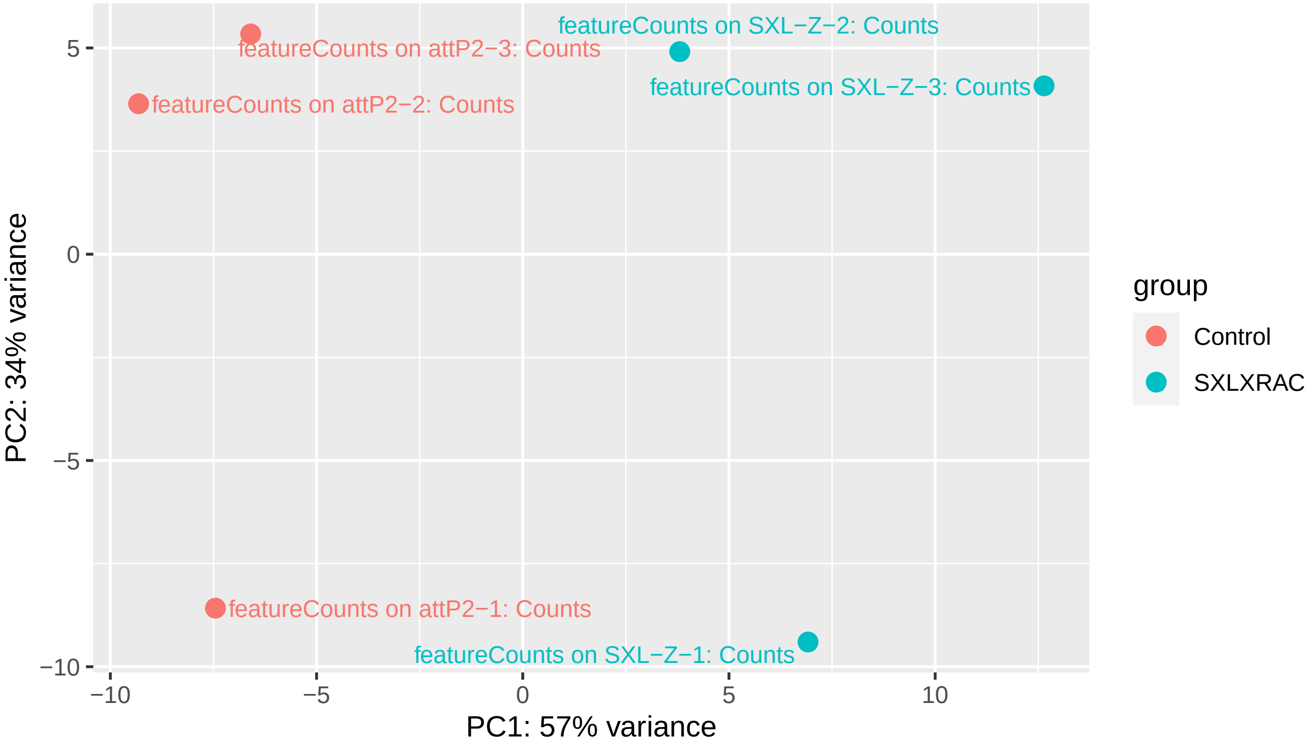

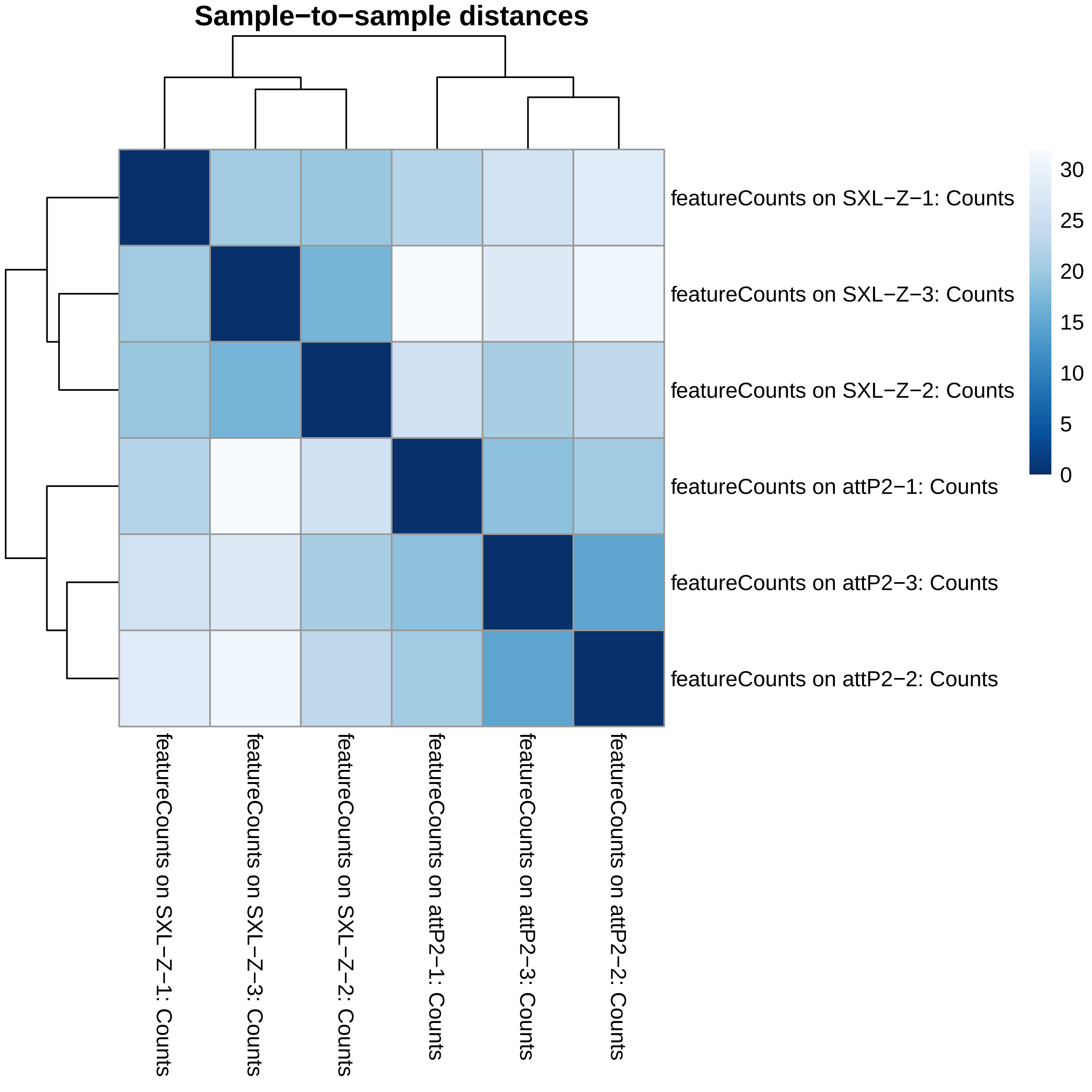

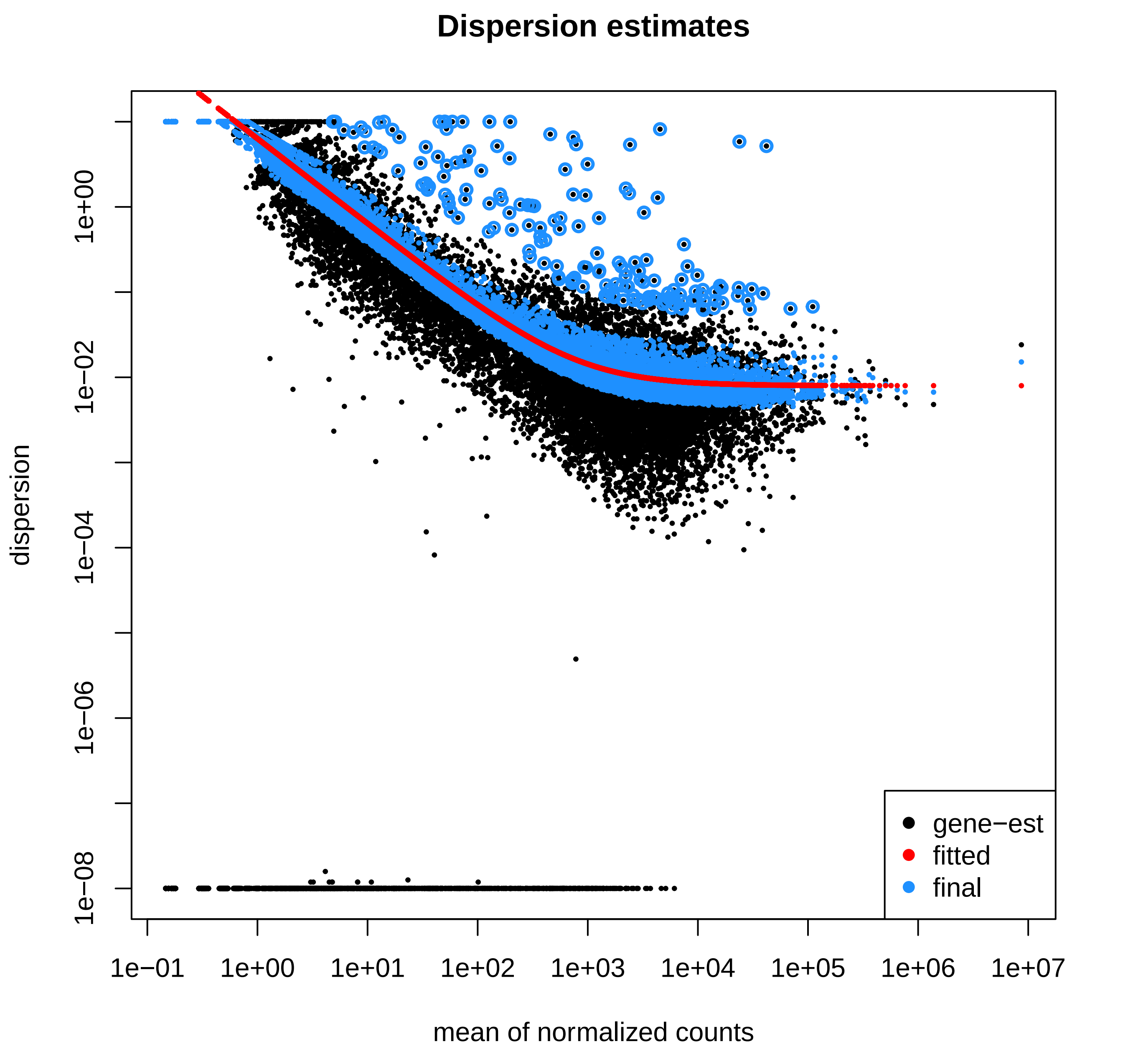

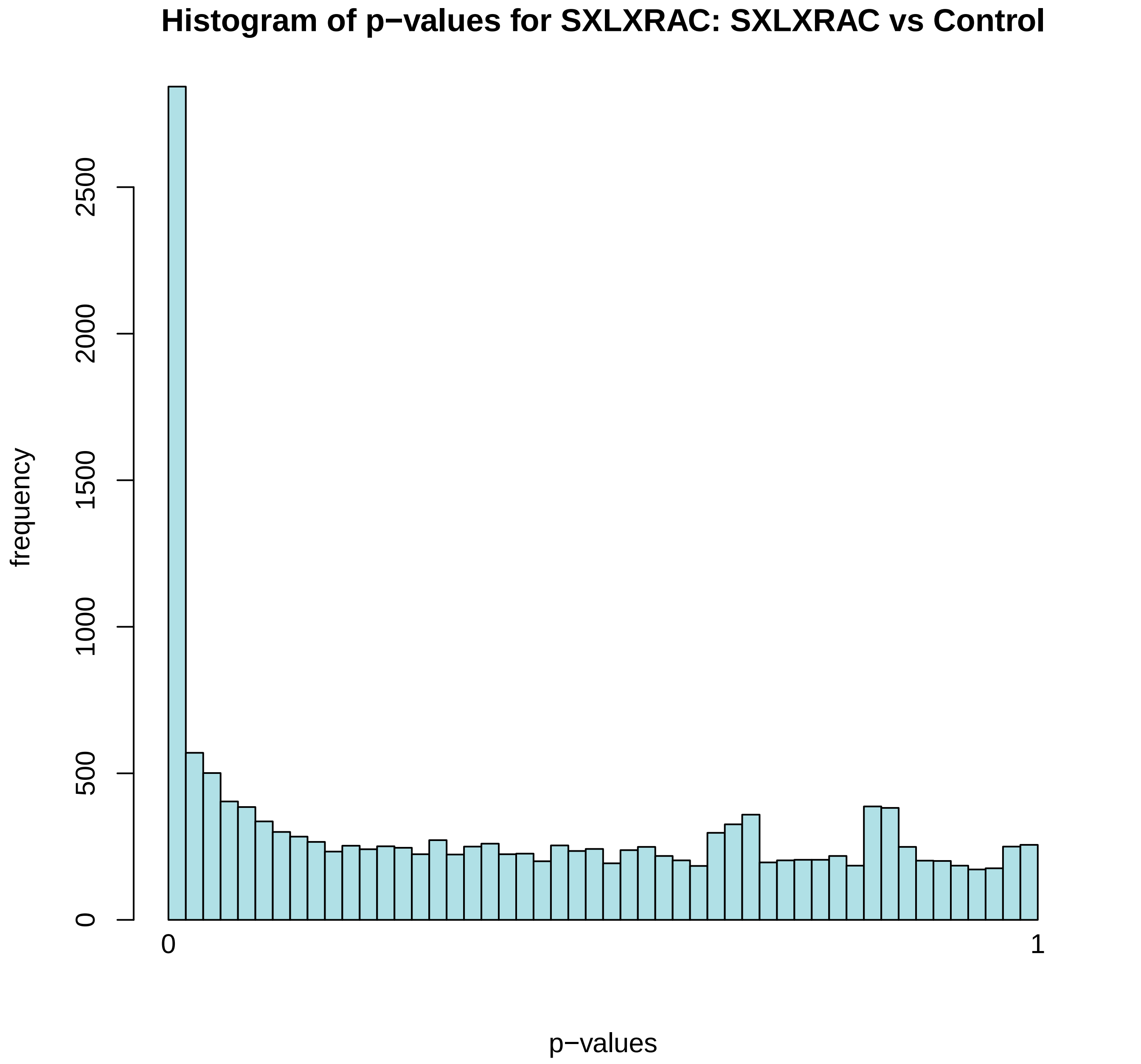

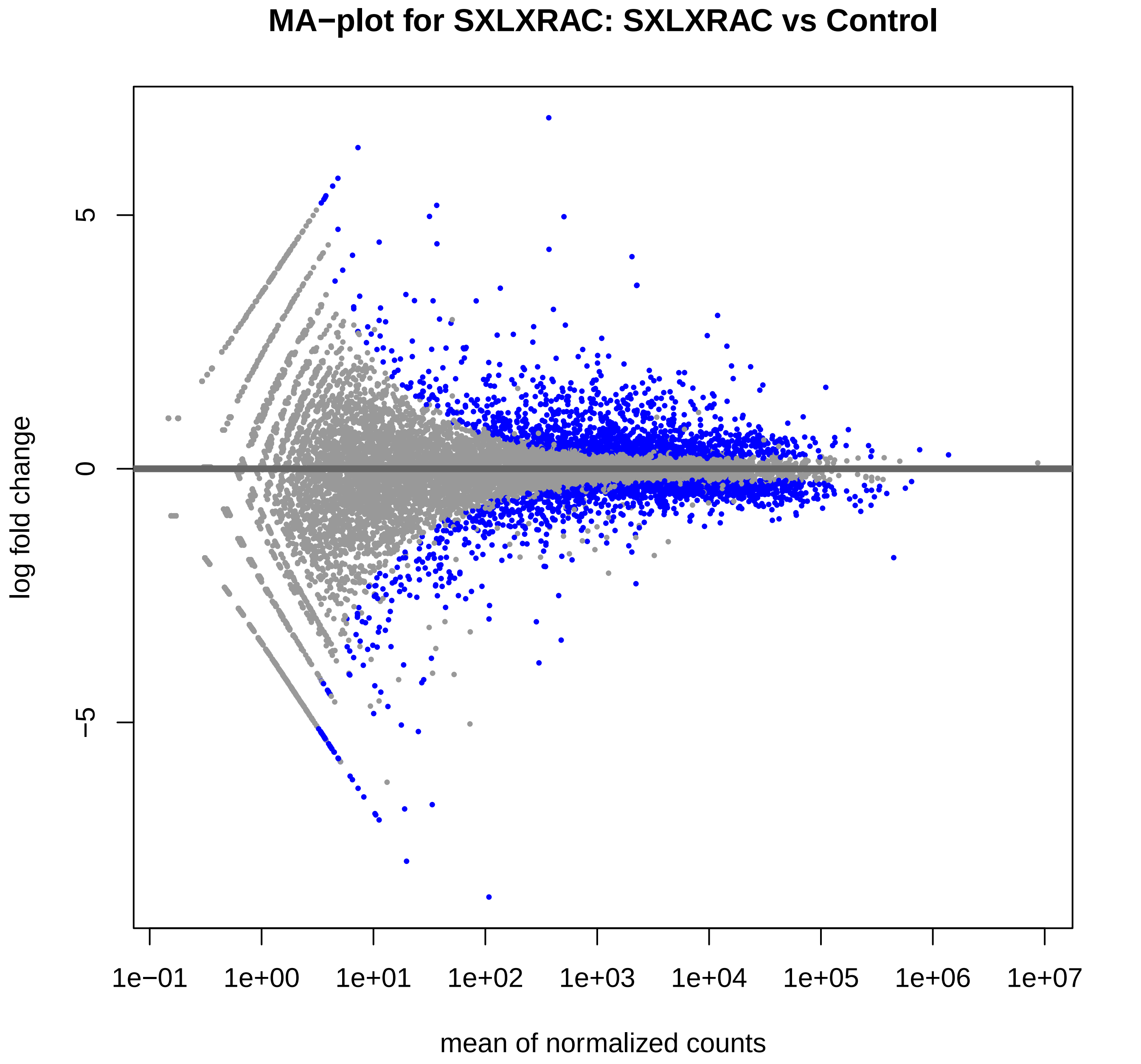

### S36_Sxl_RNA_RNAseq_Plots.pdf

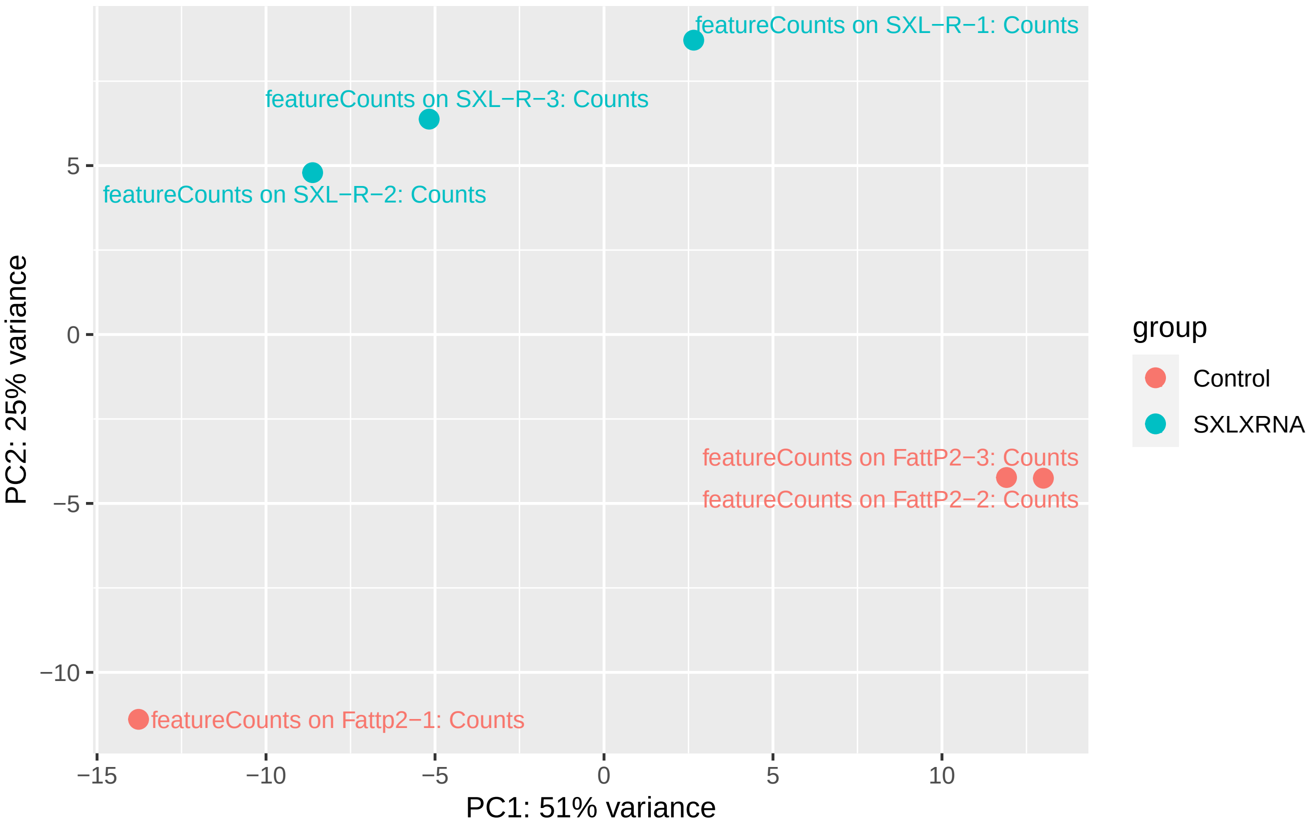

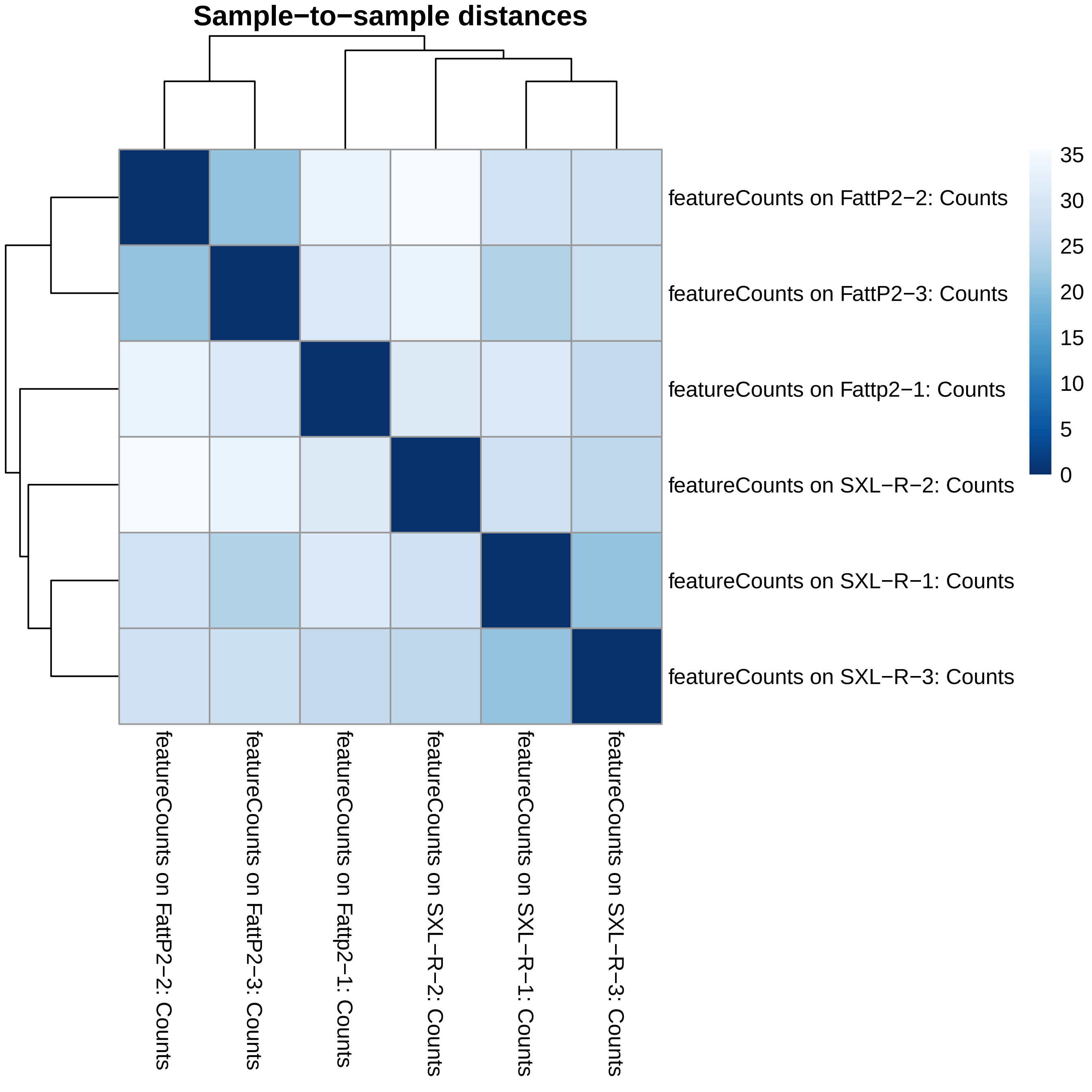

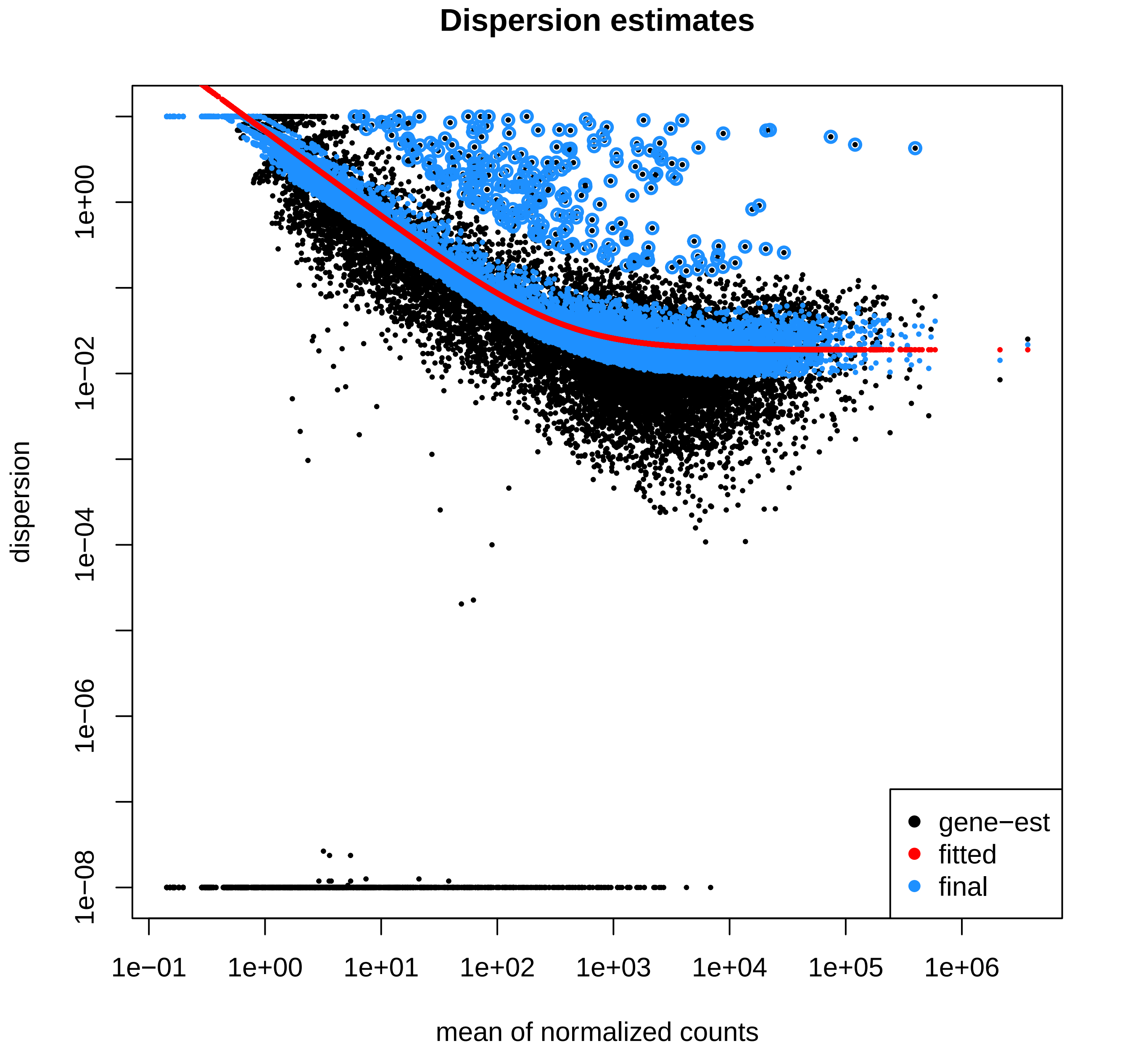

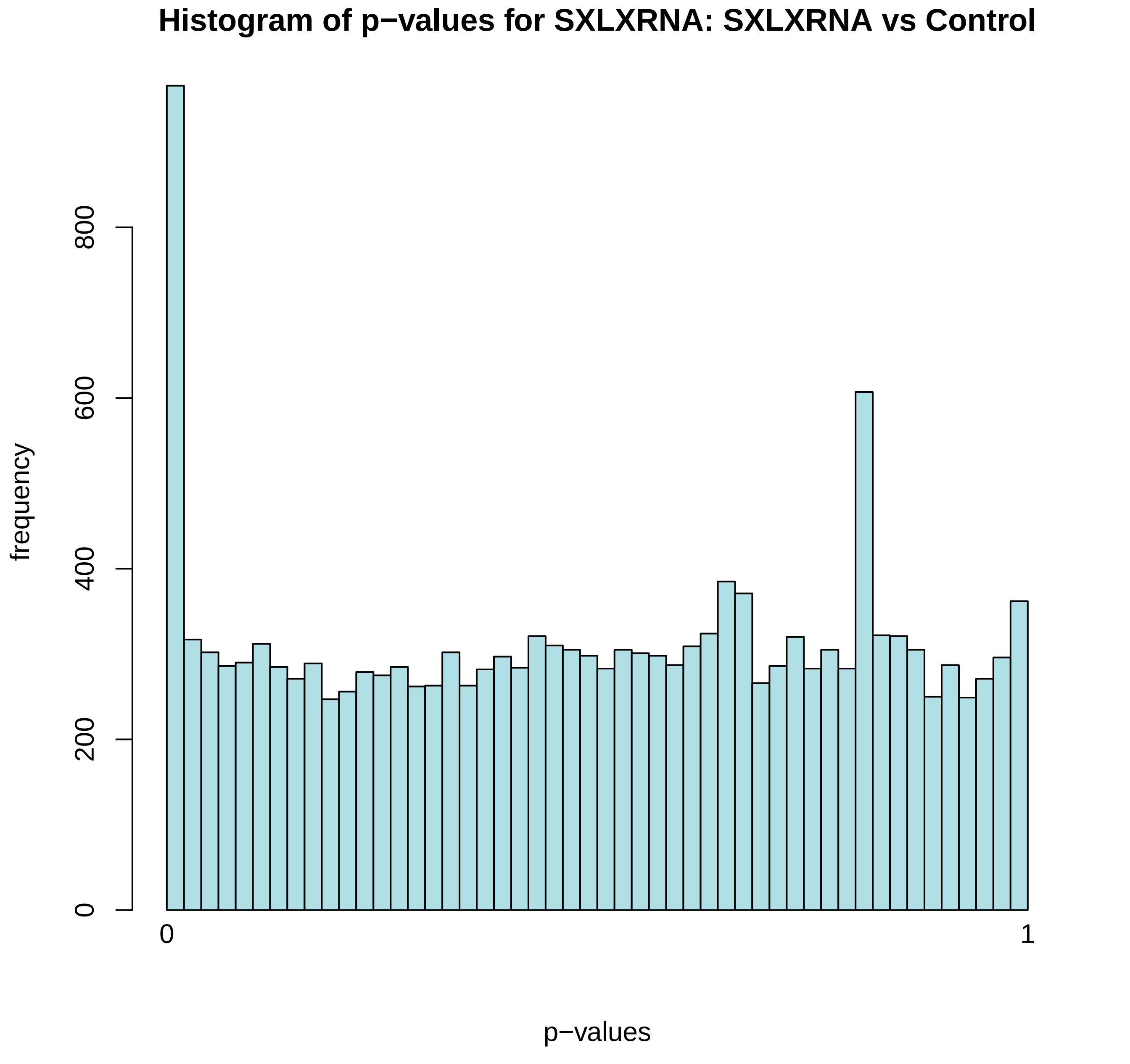

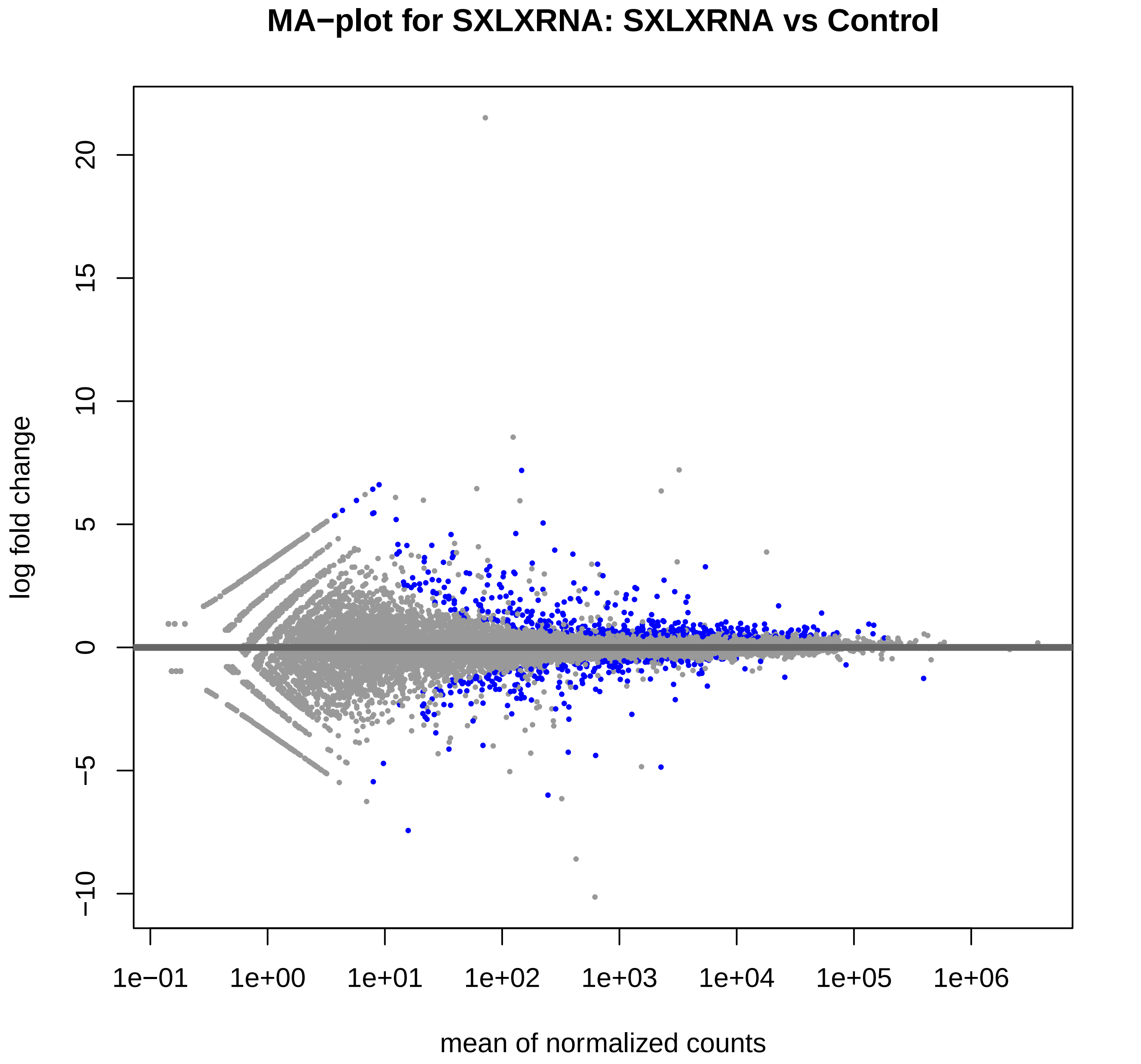
