## Supplementary material for "Sex-lethal is recruited to chromatin to promote neuronal tRNA synthesis in males through RNA Polymerase III regulation": Table of Supplementary data

S1_Sxl_DamID_peaks_and_genes

S2_Sxl_GSmut_DamID_peaks_and_genes

S3_Sxl_RNAmut_DamID_peaks_and_genes

S4_Sxl_NDel_DamID_peaks_and_genes

S5_Sxl_CDel_DamID_peaks_and_genes

S6_Polr3E_DamID_peaks_and_genes

S7_tRNA_binding_overlap_Polr3E_vs_Sxl

S8_Sxl_DamID_Binding_wt_vs_Polr3E_RNAi_GATC_fragments_in_both_datasets

S9_Sxl_RNAi_I_Survival

S10_Sxl_RNAi_I_Negative_Geotaxis

S11_Polr3E_RNAi_Survival

S12_Polr3E_RNAi_Negative_Geotaxis

S13_Sxl_RNAi_I_GO_Term_Analysis

S14_Sxl_RAC_Survival

S15_Sxl_RAC_Negative_Geotaxis

S16_Sxl_RAC_Small_RNAseq_tRNAs

S17_Sxl_RAC_Small_RNAseq_other

S18_Sxl_OPP_Colocalisation

S19_Sxl_RNAi_II_Survival

S20_Sxl_RNAi_II_Negative_Geotaxis

S21_Sxl_RNAi_I_RNAseq_p<0.05

S22_Sxl_RNAi_I_RNAseq_Plots

S23_Sxl_RNAi_I_RNAseq_Correlation_Polr3E_RNAi

S24_Polr3E_RNAi_I_RNAseq_p<0.05

S25_Polr3E_RNAi_I_RNAseq_Plots

S26_Polr3E_RNAi_I_GO_Term_Analysis

S27_Sxl_DamID_vs_Sxl_RNAi_i_RNAseq

S28_Sxl_DamID_vs_Polr3E_RNAi_i_RNAseq

S29_Sxl_RAC_RNAseq_p<0.05

S30_Sxl_RAC_RNAseq_Plots

S31_Sxl_RAC_GO_Term_Analysis

S32_Sxl_RNA_Survival

S33_Sxl_RNA_Negative_Geotaxis

S34_Sxl_RNA_male_female_eclosion_ratio

S35_Sxl_RNA_RNAseq_p<0.05

S36_Sxl_RNA_RNAseq_Plots

S37_Sxl_RNA_GO_Term_Analysis

S38_Sxl_RAC_RNAseq_Correlation_Sxl_RNAi_I

S39_Sxl_RAC_RNAseq_Correlation_Sxl_RNA

S40_Sxl_RAC_Pol_III_Targets_qPCR

S41_Sxl_RNAi_I_Pol_III_Targets_qPCR
